# HIV-1 gp120-induced lysosomal stress responses are controlled by TRPML1 redox sensors

**DOI:** 10.64898/2026.03.02.709165

**Authors:** Nirmal Kumar, Braelyn Liang, Jonathan D. Geiger

## Abstract

Increased lysosomal stress responses (LSR) are commonly implicated in the pathogenesis of neurodegenerative disorders including HIV-1-associated neurocognitive disorders (HAND). The HIV-1 envelope glycoprotein gp120 causes LSR, increases levels of ferrous iron (Fe^2+^) in the cytosol and in mitochondria, disrupts the reactive species interactome (RSI), and increases neural cell death. Here, we report that TRPML1, an endolysosome redox-sensitive cation channel, is mechanistically involved in gp120-induced neurotoxicity. TRPML1 was activated by gp120-induced increases in cytosolic reactive oxygen species (ROS) and resulted in release of Fe^2+^ from endolysosomes in levels sufficient to increase cytosolic levels of Fe^2+^ and ROS as well as decrease levels of hydrogen sulfide (H_2_S). Reduced glutathione normally buffers intracellular Fe^2+^, but gp120 decreased endolysosome glutathione levels and disrupted this regulatory control mechanism thereby promoting TRPML1-mediated Fe^2+^ efflux from endolysosomes. TRPML1 redox activation led to changes to the RSI in endolysosomes including increased ROS, lipid peroxidation, nitric oxide, and sulfane sulfur as well as decreased H_2_S. These changes were accompanied by increased cysteine oxidation of luminal proteins and endolysosome deacidification. Pharmacological inhibition of TRPML1 or knocking down expression levels of TRPML prevented these effects. Thus, our findings suggest that TRPML1 redox activation controls gp120-induced endolysosome dysfunction and iron/redox imbalance, and further implicates TRPML1 in the pathogenesis of HAND.

## Introduction

Human immunodeficiency virus-1 (HIV-1) negatively affects brain function prior to HIV-1 seroconversion [1] and causes a broad spectrum of cognitive, motor, and behavioral deficits collectively termed HIV-associated neurocognitive disorders (HAND) [2]. Antiretroviral therapy has now improved the quality of life and longevity of people living with HIV-1 (PLWH) such that PLWH are now living almost full life spans, but the prevalence of HAND has increased and it is now estimated to affect about 50% of PLWH [2,3].

The neuropathogenesis of HAND is complex and associated with HAND are chronic long-lasting neuroinflammation, increased oxidative stress, and neurodegeneration [2]. The HIV-1 envelope glycoprotein gp120 [4] continues to be implicated in the pathogenesis of HAND [5,6]; it is neurotoxic, it causes neuroinflammation, it leads to endolysosome and mitochondrial dysfunction, it increases levels of reactive oxygen species (ROS) and reactive nitrogen species (RNS), it decreases antioxidant defense activity [5–10], and it increases the oxidation of DNA, RNA, proteins, and lipids [11,12]. In addition to increases in ROS in the CNS of HAND patients [12,13] and in neurons treated with gp120 [14], other short-lived reactive molecules may also be involved in HAND [13,15].

The reactive species interactome (RSI) is a relatively new concept of cellular stress signaling that involves reactive sulfur species (RSS), RNS, ROS, and reactive carbonyl species (RCS); the RSI also includes interactions between the various reactive species and their biological targets [16,17]. Elevated levels of RCS were reported in brain tissue and cerebrospinal fluid of HAND patients [15,18–21]. Moreover, increased levels of metal ions that cause redox catastrophe in cells, including iron, have been described in HAND patients and in gp120-treated neurons [22–24].

Endolysosomes are acidic organelles well known for their physiologically and pathologically important roles they play in degradation of metabolites and cellular debris, nutrient sensing, iron storage, redox signaling [23,24], neurodegeneration and HAND [25,26]. Endolysosome alkalization (de-acidification) leads to alternations in iron levels and because endolysosomes have readily releasable stores of Fe^2+^, endolysosome dysfunction continues to be implicated in the onset of numerous pathological conditions especially those where increased levels of Fe^2+^ have been implicated [24,28–31]. Others and we have shown that gp120 disrupts endolysosome pH, Fe^2+^, and RSI homeostasis and that the release of Fe^2+^ from endolysosomes into the cytosol was sufficient to increase levels of cytosolic and mitochondrial Fe^2+^ and ROS as well as increase neural cell death [7,28,32–34].

Endolysosome-resident transient receptor potential mucolipin 1 (TRPML1) channels transport Ca^2+^, Fe^2+^, and Zn^2+^ divalent cations, and help control endolysosome homeostatic processes, autophagy, redox signaling, and oxidative stress-induced cell death [35,36]. Loss-of-function mutations in the TRPML1 gene leads to mucolipidosis type 4, and reduced TRPML1 protein expression levels or activity can cause neurodegeneration [37–39]. TRPML1 activation can also result in neurotoxic increases in cytosolic Ca^2+^, Zn^2+^, and ROS, oxidative stress-induced neuronal injury, and cell death [39–43]. Recent studies have demonstrated that TRPML1 can be directly activated by ROS and act as a redox sensor in endolysosomes [36,41]. Thus, activation of TRPML1 activity or increased protein expression levels of TRPML1, possibly through redox activation, may contribute to gp120-induced neurotoxicity and the pathogenesis of neurodegenerative disorders.

Here, we determined the extent to which redox sensing of TRPML1 channels controls gp120-induced endolysosome dysfunction and disruption of iron and redox homeostasis. We demonstrate that TRPML1 redox activation links gp120 to endolysosome Fe^2+^ release, RSI disruption, protein cysteine oxidation, and impaired endolysosome acidification. Inhibition of TRPML1 prevented these effects thus highlighting TRPML1 as a promising therapeutic target to restore iron and redox homeostasis and protect against gp120-induced endolysosome dysfunction and neurotoxicity in HAND.

## Results

Endolysosome TRPML1 channels controlled gp120-induced decreases in endolysosome Fe^2+^ and increases in cytosolic Fe^2+^. Fe^2+^ release from endolysosomes is sufficient to account for gp120-induced increases in intracellular Fe^2+^ levels [28]. To determine the involvement of endolysosome TRPML1 channels, we measured endolysosome Fe^2+^ using LysoRhonox-1 and showed that gp120 (500 pM, 4 h) significantly decreased endolysosome Fe^2+^ levels and that this effect was blocked by pretreatment with the TRPML1 inhibitors ML-SI1 (10 μM, 1 h) and Ned-19 (1 μM, 1 h) (Fig. 1A,B). gp120 significantly increased cytosolic Fe^2+^ levels measured with PhenGreen FL DA, and pretreatment with the TRPML1 inhibitors ML-SI1, Ned-19 and YM201 (5 μM, 1 h) blocked these increases (Fig. 1C). YM201 alone decreased basal cytosolic Fe^2+^ levels (Fig. 1C). As another measure of cytosolic iron accumulation, we measured protein expression levels of ferritin H (FTH1) and found that pretreatment with Ned-19 (1 μM, 1 h) significantly (*p*<0.0125) blocked gp120-induced (500 pM, 6 h) increases in FTH1 protein expression levels. For stable TRPML1 knockdown cells, TRPML1 protein expression levels were significantly decreased by about 41% (Fig. 1F), basal cytosolic Fe^2+^ levels and gp120-induced Fe^2+^ increases were significantly decreased (Fig. 1G), and FAC-induced increases in cytosolic Fe^2+^ were significantly blocked (Fig. 1H). Conversely, pretreatment with the TRPML1 activator NAADP-AM (1 μM, 1 h) potentiated gp120-induced increases in cytosolic Fe^2+^ levels (Fig. 1I).

**Fig. 1:**
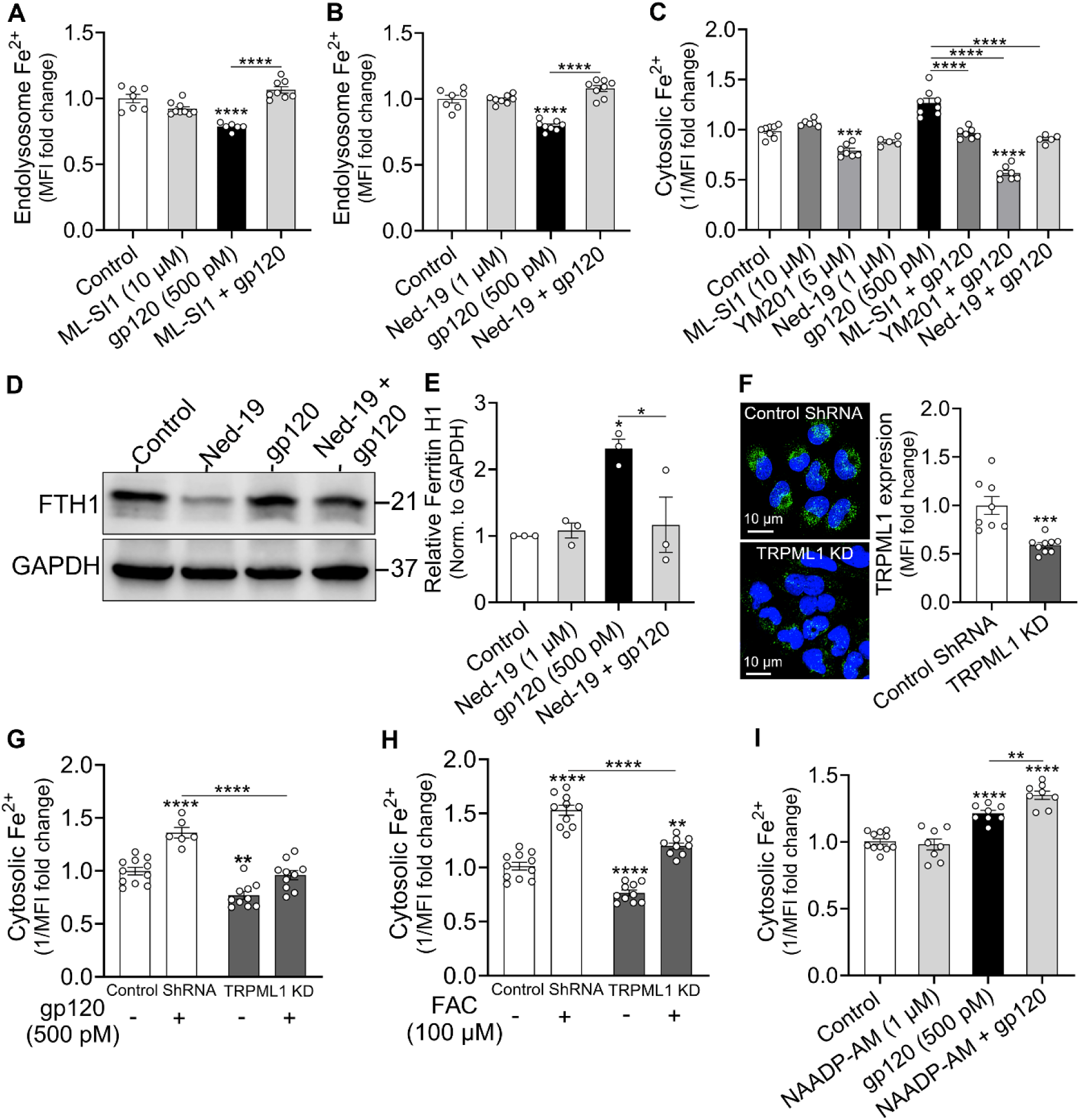
gp120-induced decreases in endolysosome Fe^2+^ and increases in cytosolic Fe^2+^ were <u>mediated by endolysosome TRPML1 channels</u> **(A, B)** Endolysosome Fe^2+^ levels were measured with LysoRhoNox-1 by flow cytometry and data were presented as fold changes of mean fluorescence intensity (MFI). gp120-induced decreases in endolysosome Fe^2+^ levels were significantly decreased by pre-treatment (1 h) with the TRPML1 inhibitors Ned-19 (**A**, 1 μM) and ML-SI1 (**B**, 10 μM). **(C)** Cytosolic Fe^2+^ levels were measured with Phen Green FL DA by flow cytometry and data were presented as reciprocals of MFI (1/MFI). gp120-induced increases in cytosolic Fe^2+^ levels were significantly decreased by the TRPML1 inhibitors ML-SI1 (10 µM), YM201 (5 µM), and Ned-19 (1 µM). YM201, but not ML-SI1 or Ned-19 significantly decreased basal cytosolic Fe^2+^ levels. **(D, E)** Representative Western blot image (**D**) and quantification of ferritin H (FTH1) protein expression levels showed that gp120-induced increases in protein expression levels of FTH1 were blocked by 1 h treatments with Ned-19 (1 µM, 1 h). **(F)** Representative immunofluorescence images of TRPML1 (green) and nuclei stained with Hoechst 33342 (blue) in U87MG cells transfected with control shRNA or TRPML1 knockdown (TRPML1 KD); TRPML1 KD significantly decreased protein expression levels of TRPML1 by ∼41%. **(G)** TRPML1 KD significantly decreased basal cytosolic Fe^2+^ levels and significantly decreased gp120-induced increases in cytosolic Fe^2+^ levels. **(H)** TRPML1 KD significantly reduced ferric ammonium citrate (FAC)-induced increases in cytosolic Fe^2+^ levels. **(I)** Pre-treatment (1 h) with the TRPML1 agonist NAADP-AM (1 µM) potentiated gp120-induced increases in cytosolic Fe^2+^ levels. Data were shown as means and SEM with individual data points (n = 4-11) included on each bar. Two-way ANOVA with Tukey’s multiple comparison tests were used for statistical analyses. \**p*<0.05, \*\**p*<0.01, \*\*\**p*<0.001, \*\*\*\**p*< 0.0001

Reduced glutathione (GSH) decreased levels of endolysosome Fe^2+^ and blocked gp120-induced increases in levels of intracellular Fe^2+^. TRPML1 is a redox sensing channel and GSH is a major anti-oxidant with metal -buffering actions [44]. Using LysoRhonox-1, we found that exogenous GSH (1 mM) but not oxidized GSH (GSSG) significantly decreased basal endolysosome Fe^2+^ levels (Fig. 2A). GSH in combination with gp120 (500 pM) further decreased endolysosome Fe^2+^ levels (Fig. 2A). As a control, we found that neither GSH (1 mM) nor GSSG (1 mM) altered Fe^2+^ binding to LysoRhonox-1 (Supplementary Fig. 5A). Using PhenGreen FL DA, GSH (1 mM) significantly decreased basal levels of Fe^2+^ in the cytosol and significantly blocked gp120-induced increases Fe^2+^ in the cytosol; GSSG (1 mM) had no significant effect (Fig. 2B). GSH and GSSG interfered with Fe^2+^ binding to PhenGreen FL (Supplementary Fig. 5B) but not FerroOrange (Supplementary Fig. 5C). Accordingly, using FerroOrange, we found that GSH significantly decreased intracellular Fe^2+^ levels, and GSH and to a lesser extent GSSG blocked gp120-induced increases in intracellular Fe^2+^ (Fig. 2C). Inhibition of endogenous GSH biosynthesis with N-methylmaleimide (NMM, 200 μM) decreased endolysosome Fe^2+^ levels (Fig. 2D) and significantly increased cytosolic Fe^2+^ levels (Fig. 2E).

**Fig. 2:**
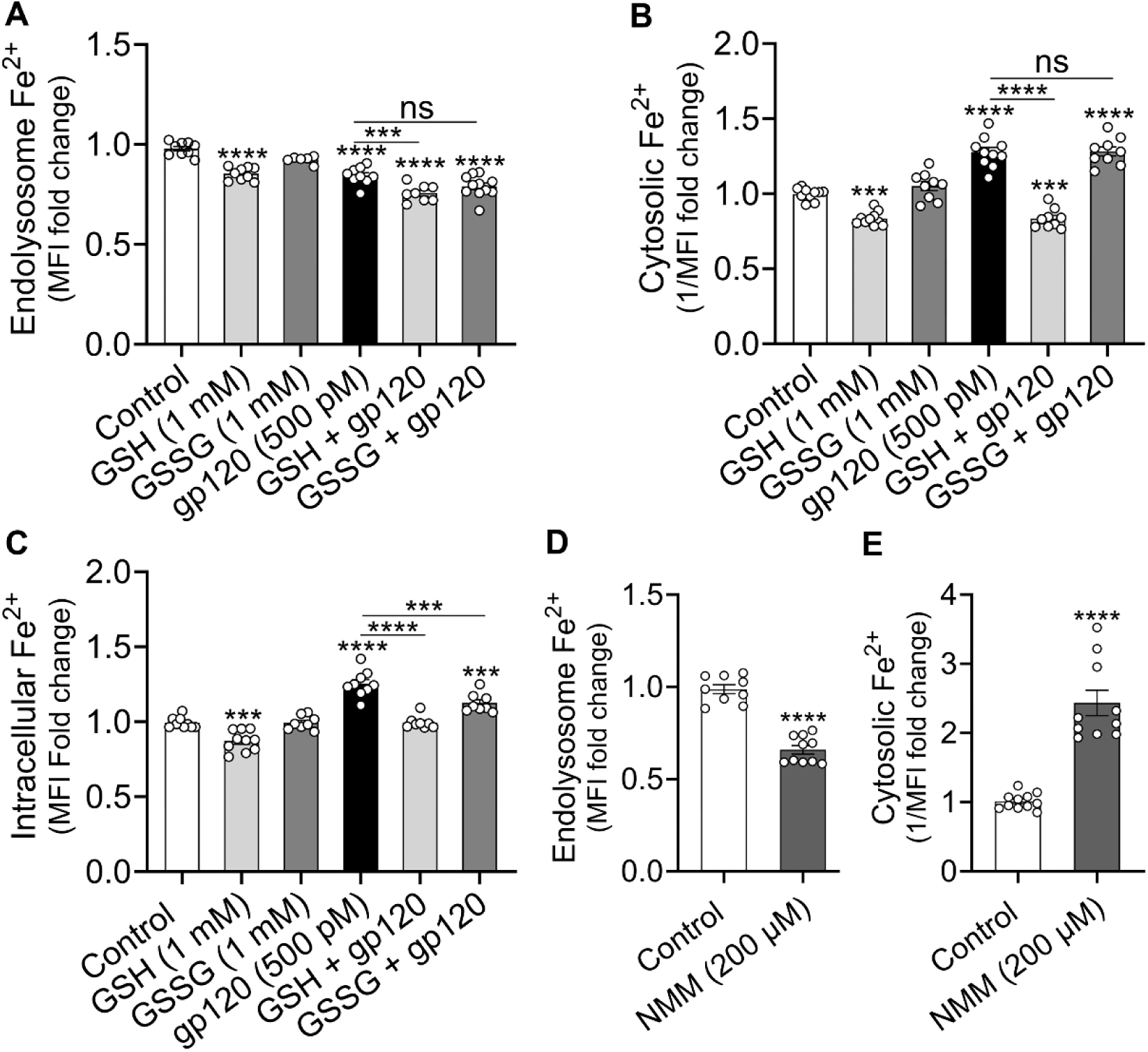
Reduced glutathione decreases levels of endolysosome and cytosolic Fe^2+^, and blocks gp120-induced decreases in endolysosome and increases in intracellular levels of Fe^2+^. **(A)** Using LysoRhoNox-1, gp120 (500 pM), GSH (1 mM), and gp120 plus GSH significantly decreased endolysosome Fe^2+^ levels. **(B)** Using Phen Green FL DA, GSH significantly decreased basal levels of cytosolic Fe^2+^ and significantly blocked gp120-induced increases in cytosolic Fe^2+^ levels. **(C)** Using FerroOrange. GSH significantly decreased basal levels of intracellular Fe^2+^, and GSH and GSSG significantly blocked gp120-induced increases in intracellular Fe^2+^ levels. **(D,E)** Inhibition of GSH biosynthesis with N-methylmaleimide (NMM, 200 µM) significantly decreased levels of endolysosome Fe^2+^ and significantly increased cytosolic Fe^2+^ levels. Data were illustrated as means and SEM with individual data points included (n = 5 to 12). Two-way ANOVA with Tukey’s multiple comparison tests were used for statistical analyses. ns = non-significant, \*\*\**p*<0.001, \*\*\*\**p*< 0.0001

gp120-induced increases in cytosolic ROS, Fe^2+^, and decreases in H_2_S were blocked by ROS <u>and TRPML1 inhibitors</u>. Using CM-H_2_DCFDA, gp120 (500 pM, 4 h) significantly increased cytosolic ROS levels, and those increases were blocked by the antioxidant NAC (5 mM) and by Ned-19 (Fig. 3A,B). As a measure of ROS, hydrogen peroxide (H_2_O_2_) concentration-dependently increased levels of cytosolic Fe^2+^ (Fig. 3C), H_2_O_2_ (100 μM)-induced increases in cytosolic Fe^2+^ were blocked by Ned-19 and ML-SI1 (Fig. 3D), and gp120-induced increases in cytosolic Fe^2+^ levels were blocked by NAC (5 mM) and Trolox (200 μM) (Fig. 3E). Using SF7-AM, gp120-induced (500 pM, 4 h) decreases in cytosolic H_2_S levels were blocked by Ned-19 (Fig. 3F). Consistent with iron involvement in H_2_S regulation [45], H_2_O_2_ (250 μM) significantly increased cytosolic ROS, serum deprivation (0% FBS) significantly increased cytosolic H_2_S levels, and iron loading with FAC (100 μM) significantly decreased H_2_S levels (Supplementary Fig. 1A,B).

**Fig. 3:**
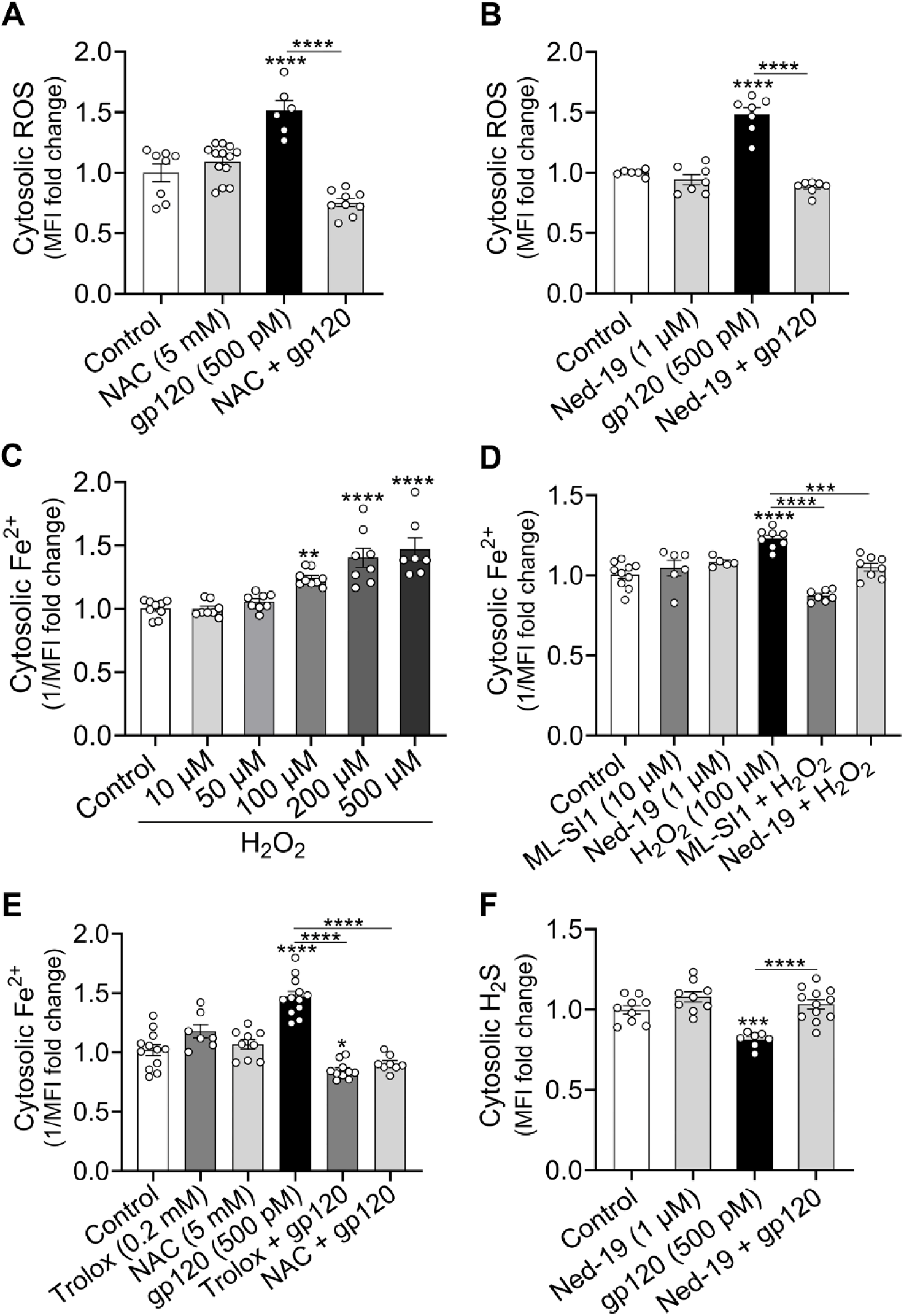
<u>gp120-induced increases in cytosolic ROS, Fe²⁺, and H_2_S were blocked by ROS and TRPML1 inhibitors</u>**. (A, B)** Using CM-H_2_DCFDA, gp120 (500 pM) significantly increased cytosolic ROS levels and those increases were significantly blocked by the antioxidant NAC (5 mM, 1 h) and the TRPML1 inhibitor Ned-19 (1 µM, 1 h). **(C,D)** Using Phen Green FL DA, H_2_O_2_ concentration-dependently increased cytosolic Fe^2+^ levels and both Ned-19 (1 µM, 1 h) and ML-SI1 (10 µM, 1 h) significantly blocked H_2_O_2-_induced increases in cytosolic Fe^2+^ levels. **(E)** The antioxidants N-acetylcysteine (NAC, 5 mM, 1 h) and Trolox (200 µM, 1 h) both significantly blocked gp120-induced increases in cytosolic Fe^2+^ levels. **(F)** Using SF7-AM, Ned-19 (1 µM, 1 h) significantly blocked gp120-induced decreases in cytosolic H_2_S levels. Data were shown as means and SEM with individual data points (n = 5 to 12). Two-way ANOVA with Tukey’s multiple comparison tests were used for statistical analyses. \**p*<0.05, \*\**p*<0.01, \*\*\**p*<0.001, \*\*\*\**p*<0.0001

<u>Ned-19 blocked gp120-induced increases in endolysosome ROS, lipid peroxidation, and nitric oxide.</u> gp120 disrupts the RSI in endolysosomes [28] and here we tested the extent to which TRPML1 redox activation mediated these effects. In LysoTracker-positive endolysosomes, gp120 (500 pM, 4 h) significantly increased CellROX (ROS), BODIPY C11(lipid peroxidation), and siRNO (nitric oxide) fluorescence (Fig. 4A-F); effects were significantly blocked by pretreatment with Ned-19 (1 μM, 1 h) (Fig. 4). gp120 activates NOX2 [46] and the NOX2 inhibitor apocynin (500 nM) significantly prevented gp120-induced increases in endolysosome ROS levels (Supplementary Fig. 2A,B). As positive controls, H_2_O_2_ (200 μM) significantly increased endolysosome ROS, RLS3 (10 μM) significantly increased endolysosome lipid peroxidation, and sodium nitroprusside (SNP, 100 μM) significantly increased endolysosome nitic oxide (Supplementary Fig. 2C-H).

**Fig. 4:**
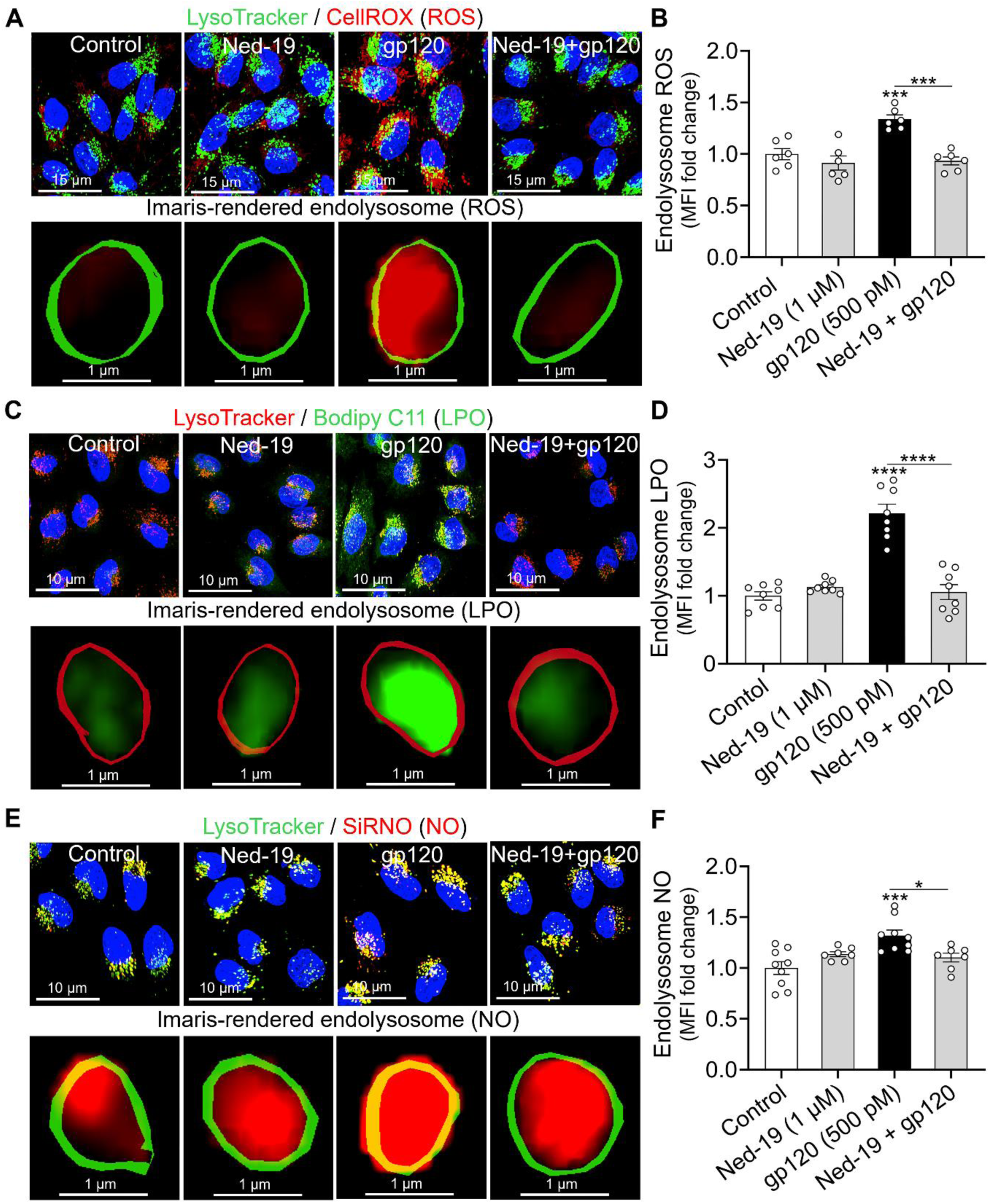
<u>Ned-19 blocked gp120-induced increases in endolysosome ROS, lipid peroxidation, and nitric oxide.</u> **(A, C, E)** Representative images of SH-SY5Y cells treated with vehicle (control), Ned-19 (1 μM, 1 h), gp120 (500 pM, 4 h) and Ned-19 plus gp120, and stained for ROS (CellROX, red), lipid peroxidation (Bodipy C11, green), nitric oxide (SiRNO, red), endolysosomes (LysoTracker, green or red), and nuclei (Hoechst 33342, blue). Scale bars were 15 µm **(A)** and 10 µm **(C, E)**. **(A, C, E)** Representative Imaris-generated surface rendering of Lysotracker-positive endolysosomes containing ROS, lipid peroxidation, and nitric oxide fluorescence. Scale bar = 1 µm. **(B, D, F)** Images were analyzed with Imaris software, and data were presented as fold-changes of mean fluorescence intensity (MFI) for CellROX, Bodipy C11 or SiRNO fluorescence within LysoTracker-positive endolysosomes. Each data point represents the fluorescence MFI within Lysotracker-positive endolysosomes from one field of view with a minimum of 100 cells analyzed per condition across at least two biological replicates. Data were shown as means and SEM with individual data points (n = 6 to 7). Two-way ANOVA with Tukey’s multiple comparison tests were used for statistical analyses. \**p*<0.05, \*\**p*<0.01, \*\*\**p*<0.001, \*\*\*\**p*<0.0001

<u>Ned-19 blocked gp120-induced decreases in endolysosome GSH and H_2_S, but not increases in H_2_S_n_.</u> gp120 (500 pM, 4 h) significantly decreased endolysosome GSH and H_2_S levels, and significantly increased sulfane sulfur (H_2_S_n_) levels (Fig. 5A-D). Ned-19 (1 μM, 1 h) significantly blocked gp120-induced decreases in GSH and H_2_S levels (Fig. 5A-D). In contrast, Ned-19 did not significantly affect gp120-induced increases in H_2_S_n_ levels (Fig. 5E,F). Ned-19 alone significantly increased endolysosome GSH, H_2_S and H_2_S_n_ levels (Fig. 5A-F). As additional validation controls, serum deprivation (0% FBS), which increases production of endogenous reactive sulfur species, significantly increased endolysosome H_2_S and H_2_S_n_ levels, and H_2_O_2_ exposure significantly decreased endolysosome GSH levels (Supplementary Fig. 3A-F).

**Fig. 5:**
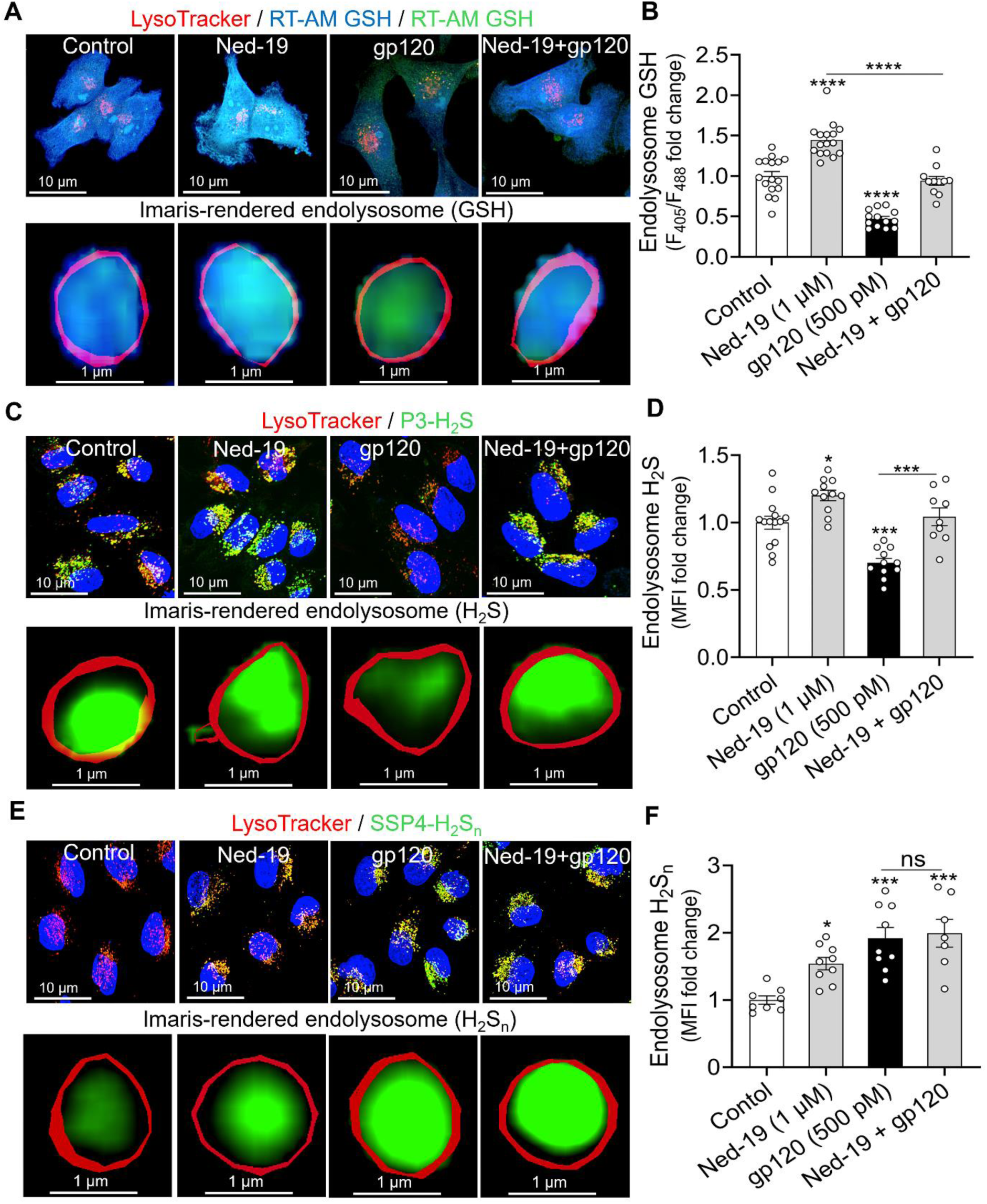
<u>Ned-19 blocked gp120-induced decreases in endolysosome GSH and H_2_S, but not increases in H_2_S_n_.</u> **(A,C,E)** Representative images of SH-SY5Y cells treated (1 h) with vehicle (control), Ned-19 (1 μM), gp120 (500 pM), and Ned-19 plus gp120. Cells were stained for GSH (RT-AM-GSH, blue/green), H_2_S (P3, green), sulfane sulfur (SSP4, green), endolysosomes (LysoTracker, red), and nuclei (Hoechst 33342, blue). Scale bars = 10 µm. **(A,C,E)** Representative Imaris-generated surface rendering of LysoTracker-positive endolysosomes containing GSH, H_2_S, or H_2_Sn. Scale bar = 1 µm. **(B,D,F)** Images were analyzed with Imaris software and data were presented as fold-changes of mean fluorescence intensity (MFI) of RT-AM GSH, P3 and SSP4 fluorescence within LysoTracker-positive endolysosomes. Each data point represents MFI within Lysotracker-positive endolysosomes from one field of view with a minimum of 100 cells analyzed per condition across at least two biological replicates. Data were shown as means and SEM with individual data points (n = 6 to 7). Two-way ANOVA with Tukey’s multiple comparison tests were used for statistical analyses. \**p*<0.05, \*\**p*<0.01, \*\*\**p*<0.001, \*\*\*\**p*<0.0001

<u>Ned-19 blocked gp120-induced increases in protein cysteine oxidation in endolysosomes.</u> Using DCP-Rho1, a dye that labels oxidized proteins at sulfenic acid sites [47], gp120 (500 pM, 1 h) significantly increased DCP-Rho1 fluorescence, Ned-19 (1 μM, 1 h) significantly decreased DCP-Rho1 fluorescence, and Ned-19 significantly blocked gp120-induced increases in DCP-Rho1 fluorescence in LysoTracker-positive endolysosomes (Fig. 6A,B). Probe specificity was confirmed by blocking DCP-Rho1 staining with dimedone and increasing DCP-Rho1 staining with H_2_O_2_ (Supplementary Fig. 4A-B).

**Fig. 6:**
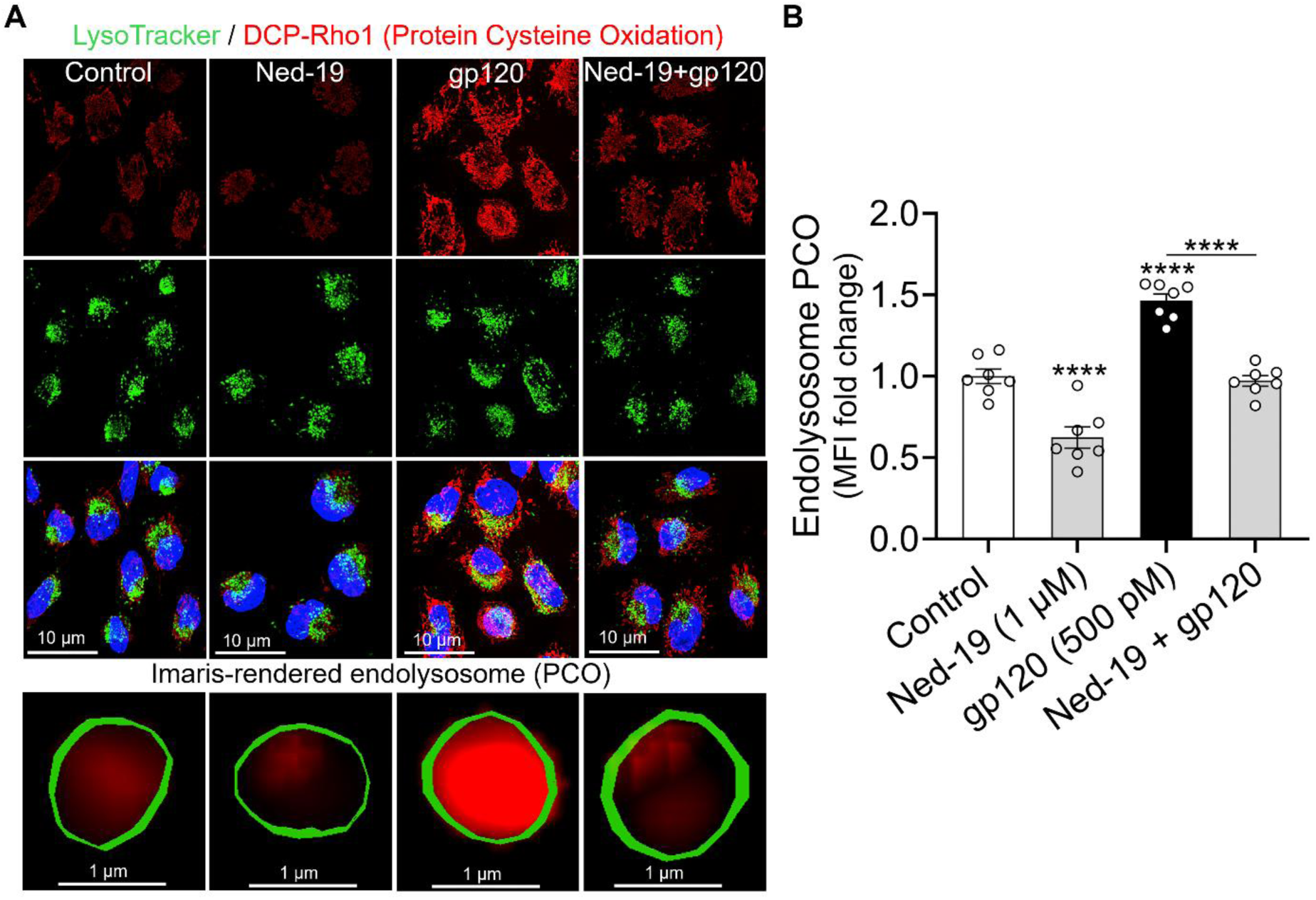
Ned-19 blocked gp120-induced increases in endolysosome protein cysteine oxidation. **(A)** Representative images of SH-SY5Y cells pre-treated for 1 h with Ned-19 (1 μM) or vehicle before treatment for 1 h with gp120 (500 pM). Cells were stained for protein cysteine oxidation (PCO; DCP-Rho-1, red), endolysosomes (LysoTracker, green) and nuclei (Hoechst 33342, blue). Scale bars = 10 µm. The bottom panels showed representative Imaris-generated surface rendering of Lysotracker-positive endolysosomes containing PCO fluorescence. Scale bar = 1 µm. **(B)** Quantification of the fold changes of mean fluorescence intensity (MFI) of DCP-Rho-1 staining within LysoTracker-positive endolysosomes showed that Ned-19 significantly decreased PCO, gp120 significantly increased PCO, and Ned-19 blocked gp120-induced increases in endolysosome PCO. Each data point represents the MFI of DCP-Rho1 staining within Lysotracker-positive endolysosomes from one field of view with a minimum of 133 cells analyzed per condition across at least two biological replicates. Data were means and SEM with individual data points (n = 7). Two-way ANOVA with Tukey’s multiple comparison tests were used for statistical analyses. \*\*\*\**p*<0.0001

<u>gp120-induced endolysosomal de-acidification was blocked by Ned-19 and antioxidants.</u> Using the ratiometric probe LysoSensor Yellow/Blue DND-160, we found that gp120 (500 pM, 30 min) significantly increased endolysosome pH (Fig. 7) and pretreatment with Ned-19 (1 μM, 30 min) blocked this effect (Fig. 7A). Because TRPML1 is a redox-sensitive channel, we next tested whether gp120-induced increases in ROS contributed to endolysosome de-acidification.

**Fig. 7:**
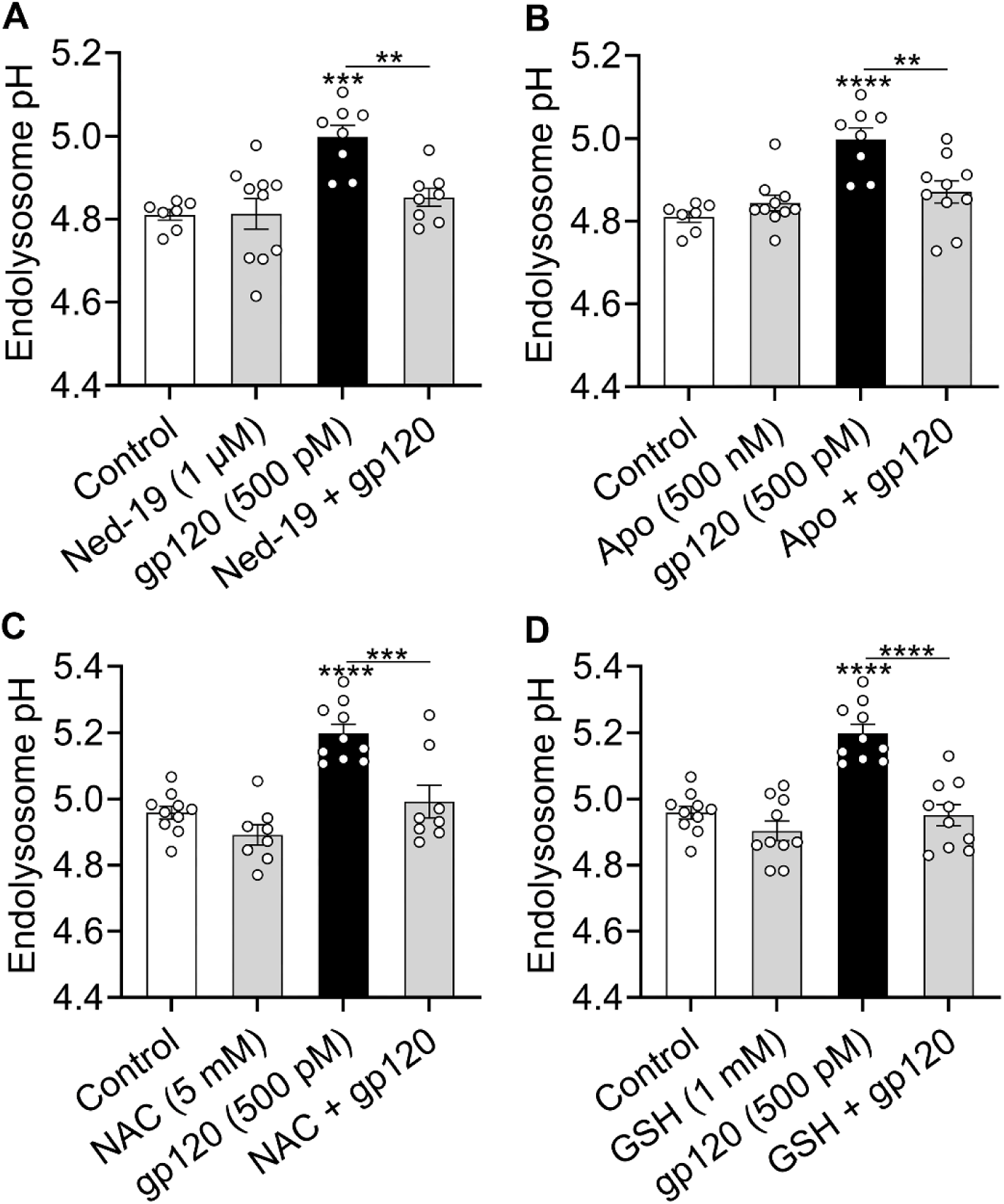
<u>Ned-19 and anti-oxidants blocked gp120-induced endolysosomal de-acidification.</u> **(A-D)** Endolysosome pH was measured in SH-SY5Y cells using LysoSensor Yellow/Blue DND-160. Cells were pre-treated for 1 h with Ned-19, apocynin, NAC, GSH or vehicle and then for 1 h with gp120 (500 pM). Each data point (n) represents the mean pH value from cells in a single well of a 96-well plate. Data were shown as means and SEM with individual data points (n = 7 to 10). Two-way ANOVA with Tukey’s multiple comparison tests were used for statistical analyses. \*\**p*<0.01, \*\*\**p*<0.001, \*\*\*\**p*<0.0001

Pretreatment with the NOX2 inhibitor apocynin (500 nM, 30 min) or the antioxidant NAC (5 mM, 30 min) prevented gp120-induced de-acidification (Fig. 7B,C). Similarly, supplementation with reduced GSH (1 mM, 30 min) also blocked gp120-induced pH changes (Fig. 7D).

## Discussion

The neurocognitive deficits that range from mild to severe with HAND have been linked to many factors including neuroinflammation and elevated levels of neurotoxic HIV-1 proteins, iron, reactive species, and organellar dysfunction [5,7,13,15,23,29]. Endolysosomes are acidic organelles that, like other organelles, exhibit stress responses to various insults; for endolysosomes this has been referred to as lysosomal stress response [48]. Others and we have shown that endolysosomes are “master regulators of cellular iron” and iron continues to be linked directly to the generation of reactive species [25,28,29,49]. Others and we have also shown that HIV-1 proteins have adverse effects on the morphology and functions of endolysosomes including increasing levels of intracellular iron, redox catastrophe, bioenergetic crisis, mitochondrial dysfunction, and increased neural cell death [24,24,24,28]. Here, we extend such work by demonstrating that reactive species once generated can feedback onto TRPML1 endolysosome cation channels that function as redox sensors and thereby induce a vicious cycle of gp120-induced iron-redox disruption, endolysosome dysfunction, inter-organellar neurotoxic signaling, and cell death. gp120-induced increases in ROS activated TRPML1 channels to increase Fe^2+^ release from endolysosomes, increase cytosolic levels of Fe^2+^ and ROS, decrease levels of cytosolic H_2_S, disrupt the endolysosome RSI, and increase endolysosome luminal protein cysteine oxidation and pH (Fig. 8).

**Fig. 8:**
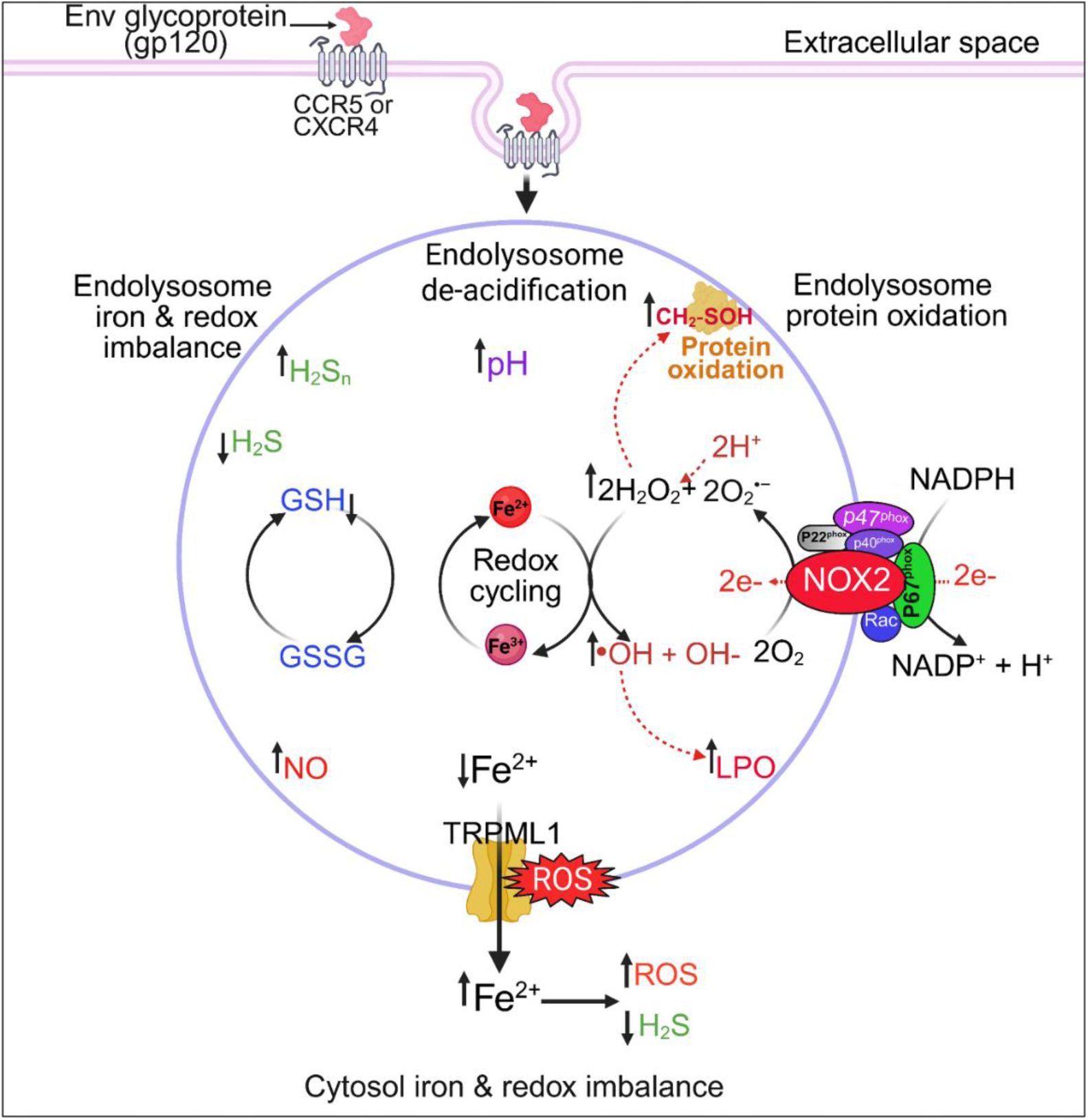
A model of gp120-induced TRPML1 redox activation and endolysosome dysfunction. gp120 is internalized and traffics to endolysosomes where it activates NOX2 and increases levels of ROS. Elevated ROS activates TRPML1, depletes endolysosome glutathione, and increases the release of Fe^2+^ into the cytosol. Increases in cytosolic Fe^2+^ and ROS and decreases in H_2_S initiate a vicious cycle of endolysosome iron release and redox disruption. TRPML1 redox activation further drives endolysosome iron and redox imbalance, impairs acidification, reduces antioxidant defenses, and induces protein cysteine oxidation and other oxidative damage. The end result is endolysosome dysfunction and increased cellular stress responses. Blocking TRPML1 channels may represent a potential therapeutic target against gp120-induced endolysosome dysfunction. Illustration was created using BioRender.

Endolysosomes contain high levels of readily releasable Fe^2+^ that are sufficient to account for insult-induced increases of Fe^2+^ and ROS in the cytosol and in mitochondria [24,28,32,49].

Insult-induced release of endolysosome Fe^2+^ occurs via multiple channels including two-pore and TRPML channels [43,50,51]. TRPML1 channels are especially important because they are activated by NAADP, the essential phosphoinositide PI(3,5)P_2_, and ROS [30,42,43]. Others and we have shown that gp120, iron overload, and other insults increase endolysosome Fe^2+^ release, increase cytosolic and mitochondrial levels of Fe^2+^ and ROS, cause mitochondrial membrane depolarization, and increase neural cell death [24,28,43,54]. Here, we showed that gp120-induced effects were TRPML1-dependent; pharmacological inhibition of TRPML channels with ML-SI1, Ned-19, and YM201636 all blocked while TRPML1 activation with NAADP-AM potentiated gp120-induced Fe^2+^ release from endolysosomes. These findings are consistent with findings that NAADP-AM enhanced whereas Ned-19 prevented TRPML1-mediated iron overload-induced cytotoxicity [43]. We also observed that FAC induced a small but statistically significant increase in cytosolic Fe^2+^ in TRPML1 knockdown cells; a finding likely mediated by TRPML2 channels as previously reported [43]. Cytosolic Fe^2+^ accumulation was accompanied by elevated ferritin expression; a cellular homeostatic mechanism that helps buffer against excess labile Fe^2+^ [55–57].

Reduced GSH, a thiol-containing redox reducing molecule, is an important endogenous anti-oxidant [44,58]. GSH and its oxidized partner GSSG protects against oxidative damage and iron dyshomeostasis intracellularly and in endolysosomes [59,60]. In neurodegenerative disorders, iron accumulation is closely linked to GSH depletion [61], as Fenton-like reactions consume GSH and promote lipid peroxidation [61]. Here, we found that exogenous GSH, but not GSSG, decreased basal levels of endolysosome Fe^2+^ as well as enhanced gp120-induced decreases in endolysosome Fe^2+^ levels. This is consistent with the ability of GSH to act as a luminal reductant thereby increasing the availability of readily releasable Fe^2+^ for TRPML1-mediated efflux [62]. Furthermore, GSH blocked gp120-induced increases in cytosolic Fe^2+^; findings consistent with reports of others that cytosolic GSH can buffer iron following its export from endolysosomes [58,58,63] and our findings that inhibition of GSH biosynthesis with NMM decreased basal endolysosome Fe^2+^ levels and increased cytosolic Fe^2+^ levels. GSH depletion elevated ferritin expression [64] and GSH regulated Zn^2+^ release from TRPM7-positive GSH-enriched organelles [65]; findings that strengthen the physiological importance of thiol-based gating of metal fluxes from acidic organelles.

TRPML1 plays dual roles in cellular signaling and homeostasis. Under physiological conditions, TRPML1 activation supports autophagy-mediated removal of damaged organelles, stimulates ROS-dependent antioxidant pathways, and promotes cell survival [36,66]. Under pathological conditions, TRPML1 activation can increase the release of endolysosome Fe^2+^, Ca^2+^, and Zn^2+^, increase levels of oxidants, disrupt autophagy and mitochondrial function, and cause neuronal death [30,34,51]. Here, we showed that gp120-induced increases in intracellular ROS activated TRPML1 to increase the release of Fe^2+^ from endolysosomes into the cytosol; effects blocked by the antioxidants NAC and Trolox. These findings are consistent with previous reports that NAC and Trolox inhibited TRPML1-mediated cell death and insult-induced increases in cytosolic levels of Zn^2+^, Ca^2+^, and ROS [41,43,66]. Further, and consistent with findings of others [30,34,51], we found that H_2_O_2_ dose-dependently increased cytosolic Fe^2+^ levels, and pharmacological inhibition of as well as knockdown of TRPML1 channels decreased H_2_O_2_-induced increases in cytosolic Fe^2+^ levels. Moreover, TRPML1 redox activation decreased levels of cytosolic H_2_S; Fe^2+^-induced increases in ROS can oxidize H_2_S to other reactive sulfur species thereby depleting H_2_S levels [67] and oxidative stress can upregulate the anti-oxidant enzyme SOD1, a potent oxidizer of H_2_S [67]. We found also that gp120-induced TRPML1 activation increased endolysosome ROS, LPO and NO levels, and decreased H_2_S, GSH and Fe^2+^ levels. These redox disruptions appeared to impair organelle function as evidenced by increased luminal protein cysteine oxidation and endolysosome de-acidification. Importantly, these effects were prevented by pretreatment with Ned-19. Thus, TRPML1 redox activation appears to link endolysosome Fe^2+^ release to a vicious self-amplifying cycle that promotes neuronal oxidative stress, mitochondrial dysfunction, and cell death.

NOX2 is a major source of ROS within endolysosomes and gp120 can activate NOX2 and increase levels of ROS and neuronal cell death [46,68,69]. Here, we found that pharmacological inhibition of NOX2 with apocynin blocked gp120-induced increases in endolysosome ROS. These results appear to be consistent with reports that HIV infection activates endolysosome NOX2 to generate ROS [70] and that TRPML1/TFEB signaling can further potential NOX2-dependent ROS production [36,71].

Hydrogen peroxide can increase oxidation of membrane phospholipids involving Fe^2+^ and Fenton-like chemical reactions [25,49,72,73]. Previously it was reported that gp120 increased LPO [28,33] and induced cortical neuronal death via increases in NO and superoxide ions [33]. Here, we found that gp120-induced increases in endolysosome LPO and NO levels.

Furthermore, increased NO production within endolysosomes led to peroxynitrite-mediated LPO increases, inhibition of vacuolar ATPase activity, and endolysosome de-acidification [74–76].

Ned-19 alone increased levels of endolysosome H_2_S, H_2_S_n_, and GSH, blocked gp120-induced decreases in H_2_S, GSH, and Fe^2+^, and restored endolysosome acidification. gp120 and HIV infection have been shown to decrease intracellular GSH and H_2_S levels [77,78]. The increased H_2_S_n_ observed here could be due to gp120-induced increases in LPO because H_2_S_n_ accumulates during LPO and contributes to LPO mitigation [79]. Importantly, H_2_S, H_2_S_n_, and GSH are synthesized by cysteine desulfuration, and together they provide protection against oxidative stress, ferroptosis, and neurodegeneration [80]. Endolysosomes are enriched in cystine, the oxidized dimeric form of cysteine, and endolysosome de-acidification has been linked to cystine depletion, lipid peroxidation, and ferroptosis [81,82]. Thus, TRPML1 redox activation-induced endolysosome de-acidification may decrease cysteine availability, decrease endolysosome H_2_S and GSH levels, and thereby exacerbate oxidative stress and cellular vulnerability to ferroptosis.

Elevated levels of reactive species can cause thiol imbalance, protein oxidation, morphological alterations, functional impairments, and neuronal cell death [44]. Endolysosomes are particularly vulnerable to oxidative damage; oxidation of v-ATPase and cathepsin B/L impairs endolysosome functions [83,84] and gp120-induced endolysosome de-acidification can decrease cathepsin B activity and cathepsin B cysteine oxidation [84]. Here, we found that Ned-19 alone decreased endolysosome protein cysteine oxidation below baseline levels, that gp120 increased endolysosome protein cysteine oxidation, and that Ned-19 blocked gp120-induced increases in endolysosome protein cysteine oxidation. Thus, TRPML1 activity can help regulate endolysosome luminal redox balance, can regulate levels of GSH, and GSH can modify protein cysteines though reversible glutathionylation, a mechanism that protects proteins from irreversible over-oxidation to sulfinic or sulfonic acids [85].

TRPML1 activation can de-acidify endolysosomes [39,40,86], Ned-19 can normalize endolysosome pH [39], and these actions are likely mediated through ROS and other reactive species production [74,83,87]. NAC restored endolysosome pH [1,2], GSH enhanced luminal enzymatic activity and acidification [90], and NOX-2-derived superoxide impaired endolysosome acidification [83,91]. Here, gp120-induced de-acidification was rescued by NAC, GSH, Ned-19, and the NOX inhibitor apocynin. Previously we showed that the endolysosome-specific iron chelator deferoxamine prevented gp120-induced endolysosome de-acidification, redox imbalance, and cell death [28,30,32,92,93]. Thus, endolysosome-specific iron chelation and inhibition of TRPML1 redox signaling may be promising approaches to protect against gp120-induced endolysosome dysfunction, iron/redox imbalance, neurotoxicity, and HAND progression.

## Materials and Methods

### Cell cultures

SH-SY5Y human neuroblastoma and U87MG human astrocytoma cells were maintained in Dulbecco’s Modified Eagle Medium (DMEM; Invitrogen, Carlsbad, USA, Cat. #11995) supplemented with 10% fetal bovine serum (FBS) and 1% penicillin/streptomycin (Invitrogen, Cat. #15140122). Cells were grown in T75 flasks and after reaching 80-90% confluency were sub-cultured using 0.025% trypsin (Invitrogen, Carlsbad USA, Cat. #25200056). All cultures were maintained in a humidified incubator at 37℃ with 5% CO_2_ and cells were not used beyond their tenth passage.

### Reagents and treatments

Recombinant HIV-1 IIIB gp120 was purchased from ImmunoDX (Woburn, MA, USA, Cat. #1001). Ferric ammonium citrate (FAC; Cat. #I72-500) was purchased from Fisher Scientific USA. Trans-Ned 19 (Cat. #3954) and NAADP tetrasodium salt (Cat. #3905) were purchased from Tocris (Minneapolis, USA). Sodium nitroprusside (SNP; Cat. #PHR1423), and sodium hydrosulfide (NaHS; Cat. #161527) were purchased from Sigma-Aldrich (St. Louis, USA). gp120 was prepared as a 0.1 µM stock solution and diluted in ultrapure distilled water to the desired concentration immediately before use. The working concentration of gp120 (500 pM; ∼60 ng/mL) is less than the reported serum levels reported in HIV-1 positive patients (120 to 960 ng/mL) [94]. Unless otherwise indicated, cells were incubated with gp120 or drugs for 1 to 4 h to investigate early effects on endolysosome function.

### Measurement of endolysosome, cytosolic and intracellular Fe^2+^, cytosolic ROS, and cytosolic H_2_S

Endolysosome Fe^2+^ levels were measured using LysoRhonox-1. Cytosolic Fe^2+^, ROS, and H_2_S were measured with Phen Green^TM^ FL diacetate (PGFL-DA), CM-H_2_DCFDA, and SF7-AM, respectively. Intracellular Fe^2+^ levels were measured with FerroOrange. The probes were purchased from the following suppliers; LysoRhonox-1 (Lumiprobe Life Science Solutions, MD, USA; Cat. #3317), PGFL-DA (Invitrogen, MA, USA; Cat. #P14313), CM-H2DCFDA (Invitrogen, Cat. #C6827), SF7-AM (Cayman Chemicals, Cat. #1416872-50-8), and FerroOrange (Dojindo, Cat. # F374). SH-SY5Y cells were seeded at 6 × 10^5^ cells/well in 24-well culture plates and incubated at 37℃ with 5% CO_2_ overnight. Cells were pre-treated with inhibitors or vehicle for 1 h followed by treatment with gp120 (500 pM) or vehicle for an additional 4 h. After treatment, cells were incubated at 37℃ with the following dyes in DMEM; 5 µM LysoRhonox-1 for 20 min, 10 µM PGFL-DA for 20 min, 10 µM CM-H2DCFDA for 20 min, 0.3 µM SF7-AM for 25 min, and 1 µM FerroOrange for 20 min. After staining, cells were washed three-times with PBS and resuspended in 600 μL PBS. Stained untreated cells served as baseline controls, and cells treated with ferric chloride (FeCl_3_, 100 μM) or ferric ammonium citrate (FAC, 50 μM) were included as positive controls for iron assays. Flow cytometry was performed on an Attune NxT flow cytometer (ThermoFisher, Waltham, USA). A minimum of 10,000 events were collected per condition. Fluorescence was recorded at the following settings; LysoRhonox-1 (Ex/Em = 561/590 nm), PLGF-DA (Ex/Em = 488/530 nm), CM-H_2_DCFDA (Ex/Em = 488/530 nm), SF7-AM (Ex/Em = 488/530 nm), and FerroOrange (Ex/Em = 561/585 nm). Each sample was measured at least three-times, and the mean MFI value was used for analyses.

### Western blot analysis of ferritin H (FTH1)

SH-SY5Y cells were seeded in 100 mm^2^ dishes and were incubated at 37℃ with 5% CO_2_ overnight. Cells were pretreated for 1 h with Ned-19 (1 μM) or vehicle, followed by treatment for 6 h with gp120 (500 pM). After treatment, cells were washed twice with ice-cold PBS and lysed in RIPA lysis buffer (ThermoScientific, Cat. #89900) supplemented with a protease inhibitor cocktail (ThermoScientific, Cat. #78429). Lysates were clarified by centrifugation (∼16000 × g for 15 min in 4℃), and protein concentrations were determined using a modified Bradford assay (ThermoScientific, Cat. #A55866). Equal amounts of protein (20 μg) were separated by SDS-PAGE on 4-12 % bis-Tris gel and were transferred onto PVDF membranes using the iBlot 3 system (ThermoFisher). Membranes were blocked in 5% non-fat milk in TBST for 1 h at room temperature and then incubated overnight at 4℃ with anti-FTH1 antibody (1:1000, Cell Signaling Technology, Cat. #4393). After washing, membranes were incubated with HRP-conjugated secondary antibody (1:3000) for 1 h at room temperature. Signals were detected using SuperSignal West Femto chemiluminescent substrate (ThermoFisher, Cat. #34095) and imaged with a ChemiDoc Imaging System (BioRad). GAPDH (1:2000) was used as a loading control. Band intensities were quantified using imageJ (Fiji) software. Western blots were performed as three independent experiments (n = 3) and quantification was based on three biological replicates.

### Measurement of endolysosome ROS, NO, lipid peroxidation, GSH, H_2_S, and sulfane sulfur

SH-SY5Y cells were seeded in 35 mm^2^ glass-bottom culture dishes and were incubated at 37℃ with 5% CO_2_ overnight. Cells were preincubated with Ned-19 (1 μM) or vehicle for 1 h followed by gp120 (500 pM) treatment for 4 h. After treatment, cells were stained in phenol red-free DMEM (Invitrogen, Cat. #21063029) with one of the following probes; CellROX^TM^ Deep Red (5 μM, Invitrogen, Cat. #C10422) for ROS (30 min), BODIPY^TM^ 581/591 C11 (10 μM, Invitrogen, Cat. #D3861) for lipid peroxidation (25 min), siRNO (5 μM, Sigma-Aldrich, Cat. #SCT053) for NO (30 min), RT-AM GSH (1 μM, Kerafast, Inc. MA USA., Cat. # EBY001) for reduced glutathione (10 min), P3 (10 μM, Sigma-Aldrich, Cat. #5.34329) for H_2_S (20 min), and SSP4 (5 μM, Dojindo, Cat. #SB10-10) for sulfane sulfur (20 min). LysoTracker^TM^ (50 nM) to label endolysosomes and Hoechst 33342 (1 μg/mL, ThermoFisher, Cat. #62249) to label nuclei were added for the final 7 min of incubations. Cells were washed three-times with PBS (Invitrogen, Cat. #14190144) and resuspended in live cell imaging solution (Invitrogen, Cat. #A59688DJ).

Imaging was performed using an Andor Dragonfly 200 spinning-disk confocal microscope (Oxford Instruments, Concord, USA) equipped with a 63x oil immersion objective and multichannel acquisition. The following microscopy settings used were as follows; CellROX (Ex/Em = 637/698 nm), siRNO (Ex/Em = 637/690 nm), P3 and SSP4 (Ex/Em = 488/530 nm), RT-AM-GSH (Ex/Em = 405 or 530 nm), BODIPY C11 (Ex/Em =488/530 nm for oxidized green and 581/600 nm for reduced red), LysoTracker deep red (647/668 nm, Invitrogen, Cat. #L12492) and LysoTracker Green (Ex/Em = 488/521 nm) for endolysosomes, and Hoechst 33342 (Ex/E = 405/445 nm) for nuclei. Mean fluorescence intensity (MFI) was analyzed using Imaris software (v9.9).

### Endolysosome protein cysteine oxidation measurements

SH-SY5Y cells were seeded in 35 mm^2^ culture dishes and incubated overnight at 37℃ and 5% CO_2_. Cells were pre-incubated with Ned-19 (1 μM) or vehicle for 1 h before addition of gp120 (500 pm) or vehicle for an additional 1 h. DCP-Rho1 (10 μM, Kerafast, Inc. MA USA., Cat. #EE0031) was added for the final 15 min, and LysoTracker^TM^ Green DND-26 (50 nM) together with Hoechst 33342 (1 μg/mL) were added for the final 7 min. Cells were washed two-times with DMEM, two-times with PBS, and then resuspended in live-cell imaging solution. Imaging was performed on an Andor Dragonfly 200 spinning disk confocal microscope equipped with 63x oil immersion objective.

Acquisition settings were as follows; Lysotracker Green (Ex/Em = 488/521 nm), DCP-Rho1 (Ex/Em = 561/590 nm), and Hoechst 33342 (Ex/Em = 405/445 nm). Images were analyzed using Imaris software (v9.9), and MFI was quantified within LysoTracker-positive endolysosomes. DCP-Rho1 selectively reacts with sulfenic acid modifications of cysteine residues and has been previously validated as a selective probe for protein cysteine oxidation [47,95]. To further confirm specificity, we pretreated with dimedone, a selective sulfenic acid scavenger to decrease DCP-Rho-1 MFI and the positive control H_2_O_2_ (100 μM) to increase DCP-Rho-1 MFI.

### Endolysosome pH measurements

Endolysosome pH was quantified using the ratiometric probe LysoSensor Yellow/Blue DND-160 (5 μM, Invitrogen, Waltham, MA, U.S. Cat. #L7545) [28]. SH-SY5Y cells were seeded in black, clear-bottom 96-well culture plates (Invitrogen, Waltham, MA, U.S. Cat. #165305) and incubated overnight at 37℃ and 5% CO_2_. Cells were pretreated with inhibitors or vehicle for 30 min followed by treatment with gp120 (500 pM) for an additional 30 min. Cell culture media was replaced with imaging buffer (140 mM NaCl, 2.5 mM KCl, 1.8 mM CaCl_2_, 1 mM MgCl_2_, and 20 mM HEPES, pH 7.40) containing 5 μM LysoSensor Yellow/Blue DND-160. Cells were incubated for 4 min at room temperature in the dark, rinsed three-times with imaging buffer to remove excess dye, and immediately analyzed. Fluorescence intensity was measured with a BioTek Synergy H1 microplate reader with dual excitation wavelengths 340 nm (F_340_) and 380 nm (F3_80_); emission was recorded at 527 nm. Relative endolysosomal pH was measured from the F_340_/F_380_ ratio and pH was quantified from calibration curves using GraphPad Prism as described previously [28].

### Effects of GSH and GSSG on Fe^2+^ detection by fluorescence probes

LysoRhoNox (5 μM, Ex/Em = 569/581), PhenGreen FL DA (5 μM, 488/530 nm), and FerroOrange (1 μM, 561/570 nm) were diluted in PBS in black clear-bottom 96-well culture plates with or without 1 mM GSH or GSSG. FeCl_2_ was then added, and fluorescence changes were measured with a BioTek Synergy H1 microplate reader.

### Generation of stable TRPML1 knockdown U87MG cells

U87MG cells were transfected with either TRPML1 shRNA plasmid (Santa Cruz, Sc-44519) or control shRNA plasmid (Sc-108060) using Jet prime reagent as per manufacturer’s instructions. After 36 h, cells were selected in puromycin (20 μg/mL; Invitrogen) for 1 week to generate stable knockdown lines. Knockdown efficiency was confirmed by immunofluorescence staining. For immunofluorescence, cells were fixed with 4% paraformaldehyde (15 min, room temperature) and permeabilized with 0.1% Triton-X 100 in PBS (10 min, room temperature). Nonspecific binding was blocked with 10% goat serum in PBS (1 h, room temperature). Cells were incubated overnight at 4℃ with anti-MCOLN1 antibody (1:500, NOVUS biologicals, CO, USA., Cat. #NBP1-92152) followed by washing and incubation with Alexa Fluor 488-conjugated secondary antibodies (1:1000) for 1 h at room temperature. Nuclei were counterstained with Hoechst 33342. Imaging was performed using our spinning disk confocal microscope with a 63x oil immersion objective. Acquisition settings included the GFP channel (Ex/Em = 488/ 521 nm) for TRPML1 and the DAPI channel (Ex/Em = 405/445 nm) for Hoechst. Images were analyzed using Imaris software (v9.9).

### Statistics and reproducibility

All experiments were independently repeated to ensure reproducibility. Comparisons between two groups were performed using Student’s *t*-test. For multiple group comparison, one-way or two-way ANOVA with Tukey’s post hoc test was used. Normality was assed using Shapiro-Wilk’s test and Kolmogorov-Smirnov’s test, and outliers were identified using the ROUT method (ROUT = 1%). A p<0.05 was considered statistically significant. All analyses were performed using GraphPad Prism (v9.4-9.5).

## Acknowledgments

This work was supported by the National Institute of General Medical Sciences [P20GM139759], the National Institute of Mental Health [R01MH119000]’, the National Institute of Neurological Disorders and Stroke [2R01NS065957], and the National Institute on Drug Abuse [2R01DA032444].

## Author contribution

N.K. conceived the study, designed and performed experiments, collected and analyzed data, interpreted results, and wrote the original draft. B.L. assisted with experiments and validation. J.D.G. contributed to conceptualization of the study, provided supervision, and edited the manuscript.

## Data and materials availability

All data supporting findings of this study are included in the manuscript and supplementary materials. Additional information is available from the corresponding author upon reasonable request.

## Conflict of Interest

None.

## Notes

### Competing Interest Statement

The authors have declared no competing interest.

## References

[1] An SF, Groves M, Gray F, et al. Early Entry and Widespread Cellular Involvement of HIV-1 DNA in Brains of HIV-1 Positive Asymptomatic Individuals. Journal of Neuropathology & Experimental Neurology. 1999;58(11):1156–1162.

[2] Gray F, Adle-Biassette H, Chretien F, et al. Neuropathology and neurodegeneration in human immunodeficiency virus infection. Pathogenesis of HIV-induced lesions of the brain, correlations with HIV-associated disorders and modifications according to treatments. Clin Neuropathol. 2001;20(4):146–155.

[3] Underwood J, De Francesco D, Cole JH, et al. Validation of a Novel Multivariate Method of Defining HIV-Associated Cognitive Impairment. Open Forum Infectious Diseases. 2019;6(6):ofz198.

[4] Capó-Vélez CM, Morales-Vargas B, García-González A, et al. The alpha7-nicotinic receptor contributes to gp120-induced neurotoxicity: implications in HIV-associated neurocognitive disorders. Sci Rep. 2018;8(1):1829.

[5] Kanmogne GD, Kennedy RC, Grammas P. HIV-1 gp120 Proteins and gp160 Peptides Are Toxic to Brain Endothelial Cells and Neurons: Possible Pathway for HIV Entry into the Brain and HIV-Associated Dementia. J Neuropathol Exp Neurol. 2002;61(11):992–1000.

[6] He X, Yang W, Zeng Z, et al. NLRP3-dependent pyroptosis is required for HIV-1 gp120-induced neuropathology. Cell Mol Immunol. 2020;17(3):283–299.

[7] Agrawal L, Louboutin J-P, Marusich E, et al. Dopaminergic neurotoxicity of HIV-1 gp120: Reactive oxygen species as signaling intermediates. Brain Research. 2010;1306:116–130.

[8] Meucci O, Miller RJ. gp120-Induced Neurotoxicity in Hippocampal Pyramidal Neuron Cultures: Protective Action of TGF-β1. J Neurosci. 1996;16(13):4080–4088.

[9] Zhang X, Green MV, Thayer SA. HIV gp120-induced neuroinflammation potentiates NMDA receptors to overcome basal suppression of inhibitory synapses by p38 MAPK. J Neurochem. 2019;148(4):499–515.

[10] Foga IO, Nath A, Hasinoff BB, et al. Antioxidants and dipyridamole inhibit HIV-1 gp120-induced free radical-based oxidative damage to human monocytoid cells. J Acquir Immune Defic Syndr Hum Retrovirol. 1997;16(4):223–229.

[11] Ellis RJ, Moore DJ, Sundermann EE, et al. Nucleic acid oxidation is associated with biomarkers of neurodegeneration in CSF in people with HIV. Neurology Neuroimmunology & Neuroinflammation. 2020;7(6):e902.

[12] Zhang Y, Wang M, Li H, et al. Accumulation of nuclear and mitochondrial DNA damage in the frontal cortex cells of patients with HIV-associated neurocognitive disorders. Brain Research. 2012;1458:1–11.

[13] Spooner RK, Taylor BK, Moshfegh CM, et al. Neuroinflammatory profiles regulated by the redox environment predicted cognitive dysfunction in people living with HIV: A cross-sectional study. eBioMedicine [Internet]. 2021 [cited 2024 May 1];70.

[14] Jana A, Pahan K. Human Immunodeficiency Virus Type 1 gp120 Induces Apoptosis in Human Primary Neurons through Redox-Regulated Activation of Neutral Sphingomyelinase. J Neurosci. 2004;24(43):9531–9540.

[15] Turchan J, Pocernich CB, Gairola C, et al. Oxidative stress in HIV demented patients and protection ex vivo with novel antioxidants. Neurology. 2003;60(2):307–314.

[16] Malard E, Valable S, Bernaudin M, et al. The Reactive Species Interactome in the Brain. Antioxidants & Redox Signaling. 2021;35(14):1176–1206.

[17] Cortese-Krott MM, Koning A, Kuhnle GGC, et al. The Reactive Species Interactome: Evolutionary Emergence, Biological Significance, and Opportunities for Redox Metabolomics and Personalized Medicine. Antioxid Redox Signal. 2017;27(10):684–712.

[18] Fennema-Notestine C, Thornton-Wells TA, Hulgan T, et al. Iron-Regulatory Genes are Associated with Neuroimaging Measures in HIV Infection. Brain Imaging Behav. 2020;14(5):2037–2049.

[19] Kaur H, Bush WS, Letendre SL, et al. Cerebrospinal Fluid Iron Status Predicts Neurocognitive Performance Over Time in Adults with HIV [Internet]. medRxiv; 2020 [cited 2024 Apr 19]. p. 2020.11.04.20199760. Available from: https://www.medrxiv.org/content/10.1101/2020.11.04.20199760v1.

[20] Cutler RG, Haughey NJ, Tammara A, et al. Dysregulation of sphingolipid and sterol metabolism by ApoE4 in HIV dementia. Neurology. 2004;63(4):626–630.

[21] Haughey NJ, Cutler RG, Tamara A, et al. Perturbation of sphingolipid metabolism and ceramide production in HIV-dementia. Annals of Neurology. 2004;55(2):257–267.

[22] Shah S, Maric D, Denaro F, et al. Nitrosative Stress Is Associated with Dopaminergic Dysfunction in the HIV-1 Transgenic Rat. Am J Pathol. 2019;189(7):1375–1385.

[23] Banjoko SO, Oseni FA, Togun RA, et al. Iron status in HIV-1 infection: implications in disease pathology. BMC Clin Pathol. 2012;12:26.

[24] Halcrow PW, Lakpa KL, Khan N, et al. HIV-1 gp120-Induced Endolysosome de-Acidification Leads to Efflux of Endolysosome Iron, and Increases in Mitochondrial Iron and Reactive Oxygen Species. J Neuroimmune Pharmacol [Internet]. 2021 [cited 2022 Feb 4]; doi: 10.1007/s11481-021-09995-2.

[25] Rizzollo F, More S, Vangheluwe P, et al. The lysosome as a master regulator of iron metabolism. Trends in Biochemical Sciences. 2021;46(12):960–975.

[26] Wang W, Gao Q, Yang M, et al. Up-regulation of lysosomal TRPML1 channels is essential for lysosomal adaptation to nutrient starvation. Proc Natl Acad Sci U S A. 2015;112(11):E1373–E1381.

[27] Herman M, Randall GW, Spiegel JL, et al. Endo-lysosomal dysfunction in neurodegenerative diseases: opinion on current progress and future direction in the use of exosomes as biomarkers. Philos Trans R Soc Lond B Biol Sci. 2024;379(1899):20220387.

[28] Kumar N, Halcrow PW, Quansah DNK, et al. Involvement of endolysosome iron in HIV-1 gp120-, morphine-, and iron supplementation-induced disruption of the reactive species interactome and induction of neurotoxicity. Redox Report [Internet]. 2025 [cited 2025 Sept 30];

[29] Halcrow PW, Kumar N, Quansah DNK, et al. Chapter 5 - Endolysosome iron: Early and upstream subcellular organellar events associated with HAND. In: Hu G, Xiong H, Buch S, editors. HIV-Associated Neurocognitive Disorders [Internet]. Academic Press; 2024 [cited 2024 Apr 19]. p. 69–79. Available from: https://www.sciencedirect.com/science/article/pii/B9780323997447000262.

[30] Halcrow PW, Kumar N, Quansah DNK, et al. Endolysosome Iron Chelation Inhibits HIV-1 Protein-Induced Endolysosome De-Acidification-Induced Increases in Mitochondrial Fragmentation, Mitophagy, and Cell Death. Cells. 2022;11(11):1811.

[31] Yambire KF, Rostosky C, Watanabe T, et al. Impaired lysosomal acidification triggers iron deficiency and inflammation in vivo. eLife. 8:e51031.

[32] Halcrow PW, Kumar N, Hao E, et al. Mu opioid receptor-mediated release of endolysosome iron increases levels of mitochondrial iron, reactive oxygen species, and cell death. NeuroImmune Pharmacology and Therapeutics [Internet]. 2022 [cited 2022 Dec 16]; doi: 10.1515/nipt-2022-0013.

[33] Corasaniti MT, Navarra M, Nisticò S, et al. Requirement for Membrane Lipid Peroxidation in HIV-1 gp120-Induced Neuroblastoma Cell Death. Biochemical and Biophysical Research Communications. 1998;246(3):686–689.

[34] Hu S, Sheng WS, Lokensgard JR, et al. Preferential sensitivity of human dopaminergic neurons to gp120-induced oxidative damage. Journal of Neurovirology. 2009;15(5–6):401–410.

[35] Cao Q, Yang Y, Zhong XZ, et al. The lysosomal Ca2+ release channel TRPML1 regulates lysosome size by activating calmodulin. J Biol Chem. 2017;292(20):8424–8435.

[36] Zhang X, Cheng X, Yu L, et al. MCOLN1 is a ROS sensor in lysosomes that regulates autophagy. Nat Commun. 2016;7(1):12109.

[37] Santoni G, Maggi F, Amantini C, et al. Pathophysiological Role of Transient Receptor Potential Mucolipin Channel 1 in Calcium-Mediated Stress-Induced Neurodegenerative Diseases. Front Physiol [Internet]. 2020 [cited 2026 Feb 28];11.

[38] Sun M, Goldin E, Stahl S, et al. Mucolipidosis type IV is caused by mutations in a gene encoding a novel transient receptor potential channel. Human Molecular Genetics. 2000;9(17):2471–2478.

[39] Lie PPY, Yoo L, Goulbourne CN, et al. Axonal transport of late endosomes and amphisomes is selectively modulated by local Ca2+ efflux and disrupted by PSEN1 loss of function. Science Advances. 2022;8(17):eabj5716.

[40] Lee J-H, McBrayer MK, Wolfe DM, et al. Presenilin 1 Maintains Lysosomal Ca(2+) Homeostasis via TRPML1 by Regulating vATPase-Mediated Lysosome Acidification. Cell Rep. 2015;12(9):1430–1444.

[41] Cheng X, Liang J, Wu D, et al. Blunting ROS/TRPML1 pathway protects AFB1-induced porcine intestinal epithelial cells apoptosis by restoring impaired autophagic flux. Ecotoxicology and Environmental Safety. 2023;257:114942.

[42] Liu Y, Wang X, Zhu W, et al. TRPML1-induced autophagy inhibition triggers mitochondrial mediated apoptosis. Cancer Lett. 2022;541:215752.

[43] Fernández B, Olmedo P, Gil F, et al. Iron-induced cytotoxicity mediated by endolysosomal TRPML1 channels is reverted by TFEB. Cell Death Dis. 2022;13(12):1–11.

[44] Glutathione, oxidative stress and neurodegeneration - Schulz - 2000 - European Journal of Biochemistry - Wiley Online Library [Internet]. [cited 2024 May 2]. Available from: https://febs.onlinelibrary.wiley.com/doi/full/10.1046/j.1432-1327.2000.01595.x.

[45] Arif HM, Qian Z, Wang R. Signaling Integration of Hydrogen Sulfide and Iron on Cellular Functions. Antioxidants & Redox Signaling. 2022;36(4–6):275–293.

[46] Smith LK, Babcock IW, Minamide LS, et al. Direct interaction of HIV gp120 with neuronal CXCR4 and CCR5 receptors induces cofilin-actin rod pathology via a cellular prion protein- and NOX-dependent mechanism. PLOS ONE. 2021;16(3):e0248309.

[47] Klomsiri C, Rogers LC, Soito L, et al. Endosomal H2O2 production leads to localized cysteine sulfenic acid formation on proteins during lysophosphatidic acid-mediated cell signaling. Free Radic Biol Med. 2014;71:49–60.

[48] Lakpa KL, Khan N, Afghah Z, et al. Lysosomal Stress Response (LSR): Physiological Importance and Pathological Relevance. J Neuroimmune Pharmacol. 2021;16(2):219–237.

[49] Halcrow PW, Kumar N, Afghah Z, et al. Heterogeneity of ferrous iron-containing endolysosomes and effects of endolysosome iron on endolysosome numbers, sizes, and localization patterns. Journal of Neurochemistry [Internet]. [cited 2022 Mar 4];n/a(n/a).

[50] Dong X-P, Cheng X, Mills E, et al. The type IV mucolipidosis-associated protein TRPML1 is an endolysosomal iron release channel. Nature. 2008;455(7215):992–996.

[51] Jin X, Zhang Y, Alharbi A, et al. Targeting Two-Pore Channels: Current Progress and Future Challenges. Trends in Pharmacological Sciences. 2020;41(8):582–594.

[52] Zhang F, Jin S, Yi F, et al. TRP-ML1 functions as a lysosomal NAADP-sensitive Ca2+ release channel in coronary arterial myocytes. J Cell Mol Med. 2009;13(9B):3174–3185.

[53] Gan N, Han Y, Zeng W, et al. Structural mechanism of allosteric activation of TRPML1 by PI(3,5)P2 and rapamycin. Proc Natl Acad Sci U S A. 2022;119(7):e2120404119.

[54] Hu S, Sheng WS, Lokensgard JR, et al. Preferential sensitivity of human dopaminergic neurons to gp120-induced oxidative damage. J Neurovirol. 2009;15(5–6):401–410.

[55] Arosio P, Levi S. Cytosolic and mitochondrial ferritins in the regulation of cellular iron homeostasis and oxidative damage. Biochimica et Biophysica Acta (BBA) - General Subjects. 2010;1800(8):783–792.

[56] Garner B, Li W, Roberg K, et al. On the Cytoprotective Role of Ferritin in Macrophages and its Ability to Enhance Lysosomal Stability. Free Radical Research. 1997;27(5):487–500.

[57] Garner B, Roberg K, Brunk UT. Endogenous ferritin protects cells with iron-laden lysosomes against oxidative stress. Free Radical Research. 1998;29(2):103–114.

[58] Berndt C, Lillig CH. Glutathione, Glutaredoxins, and Iron. Antioxid Redox Signal. 2017;27(15):1235–1251.

[59] Glutathione, Glutaredoxins, and Iron | Antioxidants & Redox Signaling [Internet]. [cited 2024 May 2]. Available from: https://www.liebertpub.com/doi/abs/10.1089/ars.2017.7132.

[60] Lockwood TD. Lysosomal metal, redox and proton cycles influencing the CysHis cathepsin reaction. Metallomics. 2013;5(2):110–124.

[61] Bertrand RL. Iron accumulation, glutathione depletion, and lipid peroxidation must occur simultaneously during ferroptosis and are mutually amplifying events. Med Hypotheses. 2017;101:69–74.

[62] Terman A, Kurz T. Lysosomal Iron, Iron Chelation, and Cell Death. Antioxidants & Redox Signaling. 2013;18(8):888–898.

[63] O’Keeffe R, Latunde-Dada GO, Chen Y-L, et al. Glutathione and the intracellular labile heme pool. Biometals. 2021;34(2):221–228.

[64] Morozova N, Khrapko K, Panee J, et al. Glutathione depletion in hippocampal cells increases levels of H and L ferritin and glutathione S-transferase mRNAs. Genes to Cells. 2007;12(5):561–567.

[65] Abiria SA, Krapivinsky G, Sah R, et al. TRPM7 senses oxidative stress to release Zn2+ from unique intracellular vesicles. Proceedings of the National Academy of Sciences. 2017;114(30):E6079–E6088.

[66] Tedeschi V, Sisalli MJ, Petrozziello T, et al. Lysosomal calcium is modulated by STIM1/TRPML1 interaction which participates to neuronal survival during ischemic preconditioning. FASEB J. 2021;35(2):e21277.

[67] Switzer CH, Kasamatsu S, Ihara H, et al. SOD1 is an essential H2S detoxifying enzyme. Proceedings of the National Academy of Sciences. 2023;120(3):e2205044120.

[68] Shah A, Kumar S, Simon SD, et al. HIV gp120- and methamphetamine-mediated oxidative stress induces astrocyte apoptosis via cytochrome P450 2E1. Cell Death Dis. 2013;4(10):e850–e850.

[69] Samikkannu T, Ranjith D, Rao KVK, et al. HIV-1 gp120 and morphine induced oxidative stress: role in cell cycle regulation. Frontiers in Microbiology [Internet]. 2015 [cited 2022 Feb 4];6.

[70] Endosomal NOX2 oxidase exacerbates virus pathogenicity and is a target for antiviral therapy | Nature Communications [Internet]. [cited 2024 May 24]. Available from: https://www.nature.com/articles/s41467-017-00057-x.

[71] Najibi M, Honwad HH, Moreau JA, et al. A NOVEL NOX/PHOX-CD38-NAADP-TFEB AXIS IMPORTANT FOR MACROPHAGE ACTIVATION DURING BACTERIAL PHAGOCYTOSIS. Autophagy. 2022;18(1):124–141.

[72] Kurz T, Gustafsson B, Brunk UT. Intralysosomal iron chelation protects against oxidative stress-induced cellular damage. The FEBS Journal. 2006;273(13):3106–3117.

[73] Lin Y, Epstein DL, Liton PB. Intralysosomal Iron Induces Lysosomal Membrane Permeabilization and Cathepsin D–Mediated Cell Death in Trabecular Meshwork Cells Exposed to Oxidative Stress. Investigative Ophthalmology & Visual Science. 2010;51(12):6483–6495.

[74] Qian Q, Zhang Z, Li M, et al. Hepatic Lysosomal iNOS Activity Impairs Autophagy in Obesity. Cellular and Molecular Gastroenterology and Hepatology. 2019;8(1):95–110.

[75] Bao J-X, Jin S, Zhang F, et al. Activation of Membrane NADPH Oxidase Associated with Lysosome-Targeted Acid Sphingomyelinase in Coronary Endothelial Cells. Antioxidants & Redox Signaling. 2010;12(6):703–712.

[76] Jiang L, Zheng H, Lyu Q, et al. Lysosomal nitric oxide determines transition from autophagy to ferroptosis after exposure to plasma-activated Ringer’s lactate. Redox Biology. 2021;43:101989.

[77] Pal VK, Agrawal R, Rakshit S, et al. Hydrogen sulfide blocks HIV rebound by maintaining mitochondrial bioenergetics and redox homeostasis. Kirchhoff F, Kana BD, Kirchhoff F, editors. eLife. 2021;10:e68487.

[78] Price TO, Ercal N, Nakaoke R, et al. HIV-1 viral proteins gp120 and Tat induce oxidative stress in brain endothelial cells. Brain Research. 2005;1045(1):57–63.

[79] Hydropersulfides inhibit lipid peroxidation and ferroptosis by scavenging radicals | Nature Chemical Biology [Internet]. [cited 2024 May 24]. Available from: https://www.nature.com/articles/s41589-022-01145-w.

[80] Sulfane Sulfur in Toxicology: A Novel Defense System Against Electrophilic Stress | Toxicological Sciences | Oxford Academic [Internet]. [cited 2024 May 2]. Available from: https://academic.oup.com/toxsci/article/170/1/3/5460240.

[81] Badgley MA, Kremer DM, Maurer HC, et al. Cysteine depletion induces pancreatic tumor ferroptosis in mice. Science. 2020;368(6486):85–89.

[82] Armenta DA, Laqtom NN, Alchemy G, et al. Ferroptosis inhibition by lysosome-dependent catabolism of extracellular protein. Cell Chemical Biology. 2022;29(11):1588–1600.e7.

[83] Jaishy B, Zhang Q, Chung HS, et al. Lipid-induced NOX2 activation inhibits autophagic flux by impairing lysosomal enzyme activity [S]. Journal of Lipid Research. 2015;56(3):546–561.

[84] NADPH oxidase activity controls phagosomal proteolysis in macrophages through modulation of the lumenal redox environment of phagosomes | PNAS [Internet]. [cited 2024 May 2]. Available from: https://www.pnas.org/doi/full/10.1073/pnas.0914867107.

[85] Redox Regulation via Glutaredoxin-1 and Protein S-Glutathionylation | Antioxidants & Redox Signaling [Internet]. [cited 2024 May 2]. Available from: https://www.liebertpub.com/doi/full/10.1089/ars.2019.7963.

[86] Lee C, Eldridge MJG, Gonzalez-Lozano MA, et al. DMXL1 promotes recruitment of V1-ATPase to lysosomes upon TRPML1 activation. Nat Struct Mol Biol. 2025;32(10):2060–2075.

[87] Guha S, Baltazar GC, Coffey EE, et al. Lysosomal alkalinization, lipid oxidation, and reduced phagosome clearance triggered by activation of the P2X7 receptor. The FASEB Journal. 2013;27(11):4500–4509.

[88] Song SB, Hwang ES. High Levels of ROS Impair Lysosomal Acidity and Autophagy Flux in Glucose-Deprived Fibroblasts by Activating ATM and Erk Pathways. Biomolecules. 2020;10(5):761.

[89] Tripathi A, Thangaraj A, Chivero ET, et al. N-Acetylcysteine Reverses Antiretroviral-Mediated Microglial Activation by Attenuating Autophagy-Lysosomal Dysfunction. Front Neurol [Internet]. 2020 [cited 2024 May 2];11.

[90] Fronczyk W. Effect of reduced glutathione (GSH) on activity of lysosomal system in subcellular fractions of mouse kidney.

[91] Zhan Y, Chen Q, Song Y, et al. Berbamine Hydrochloride inhibits lysosomal acidification by activating Nox2 to potentiate chemotherapy-induced apoptosis via the ROS-MAPK pathway in human lung carcinoma cells. Cell Biol Toxicol [Internet]. 2022 [cited 2023 Jan 29]; doi: 10.1007/s10565-022-09756-8.

[92] Halcrow PW, Quansah DNK, Kumar N, et al. HERV-K (HML-2) Envelope Protein Induces Mitochondrial Depolarization and Neurotoxicity via Endolysosome Iron Dyshomeostasis. J Neurosci. 2024;44(14):e0826232024.

[93] Halcrow PW, Quansah DNK, Kumar N, et al. Weak base drug-induced endolysosome iron dyshomeostasis controls the generation of reactive oxygen species, mitochondrial depolarization, and cytotoxicity. NeuroImmune Pharmacology and Therapeutics. 2024;3(1):33–46.

[94] How Much Gp120 Is There? | The Journal of Infectious Diseases | Oxford Academic [Internet]. [cited 2022 Mar 30]. Available from: https://academic.oup.com/jid/article/201/8/1273/865578?login=false.

[95] Morales Y, Nitzel DV, Price OM, et al. Redox Control of Protein Arginine Methyltransferase 1 (PRMT1) Activity. J Biol Chem. 2015;290(24):14915–14926.

