## Supplemental data for "HIV-1 gp120-induced lysosomal stress responses are controlled by TRPML1 redox sensors"

**The PDF file includes:**

Supplementary Fig.1 to 5

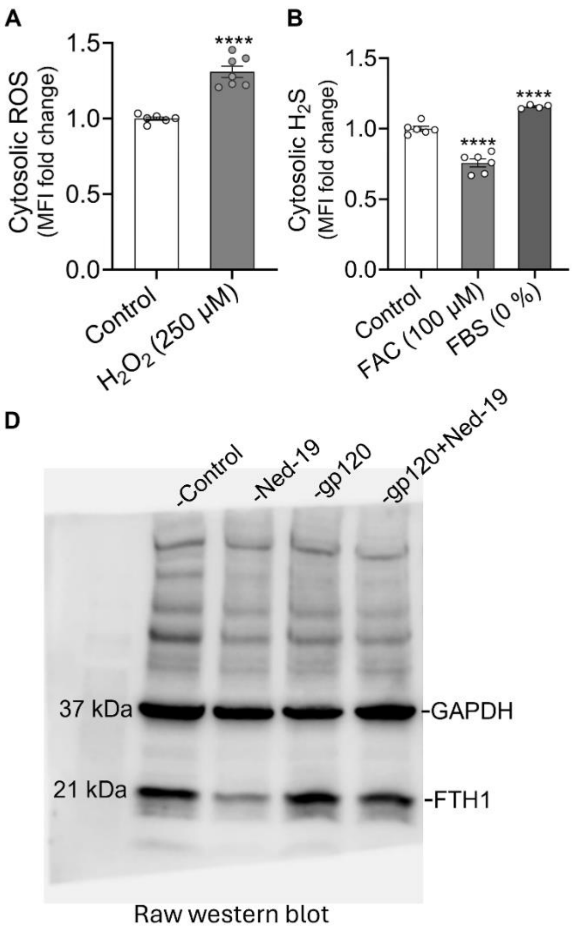


**Supplementary Fig. 1:** <u>$H_2O_2$ increased cytosolic ROS, serum deprivation increased cytosolic $H_2S$, and FAC decreased cytosolic $H_2S$.</u> **(A)** SH-SY5Y cells were treated with $H_2O_2$ (250 μM, 4 h) or vehicle control, stained with CM-$H_2$DCFDA, and fluorescence was measured by flow cytometry. $H_2O_2$ significantly increased cytosolic ROS levels. **(B)** SH-SY5Y cells were treated with FAC (100 μM, 4 h), vehicle control or incubated in FBS-free medium (0% FBS, 4 h) versus complete medium. Cells were stained with SF7-AM to detect $H_2S$ and fluorescence was measured by flow cytometry. FAC significantly decreased cytosolic $H_2S$ levels, whereas serum deprivation significantly increased $H_2S$. **(C)** Representative uncropped raw western blot image corresponding to the cropped blot shown in main Fig. 1D. Data are shown as mean ± SEM with individual data points (n = 6-7). Two-tailed Student's *t*-test or one-way ANOVA with Tukey's multiple comparison was used for statistical analysis. ****$p < 0.0001$

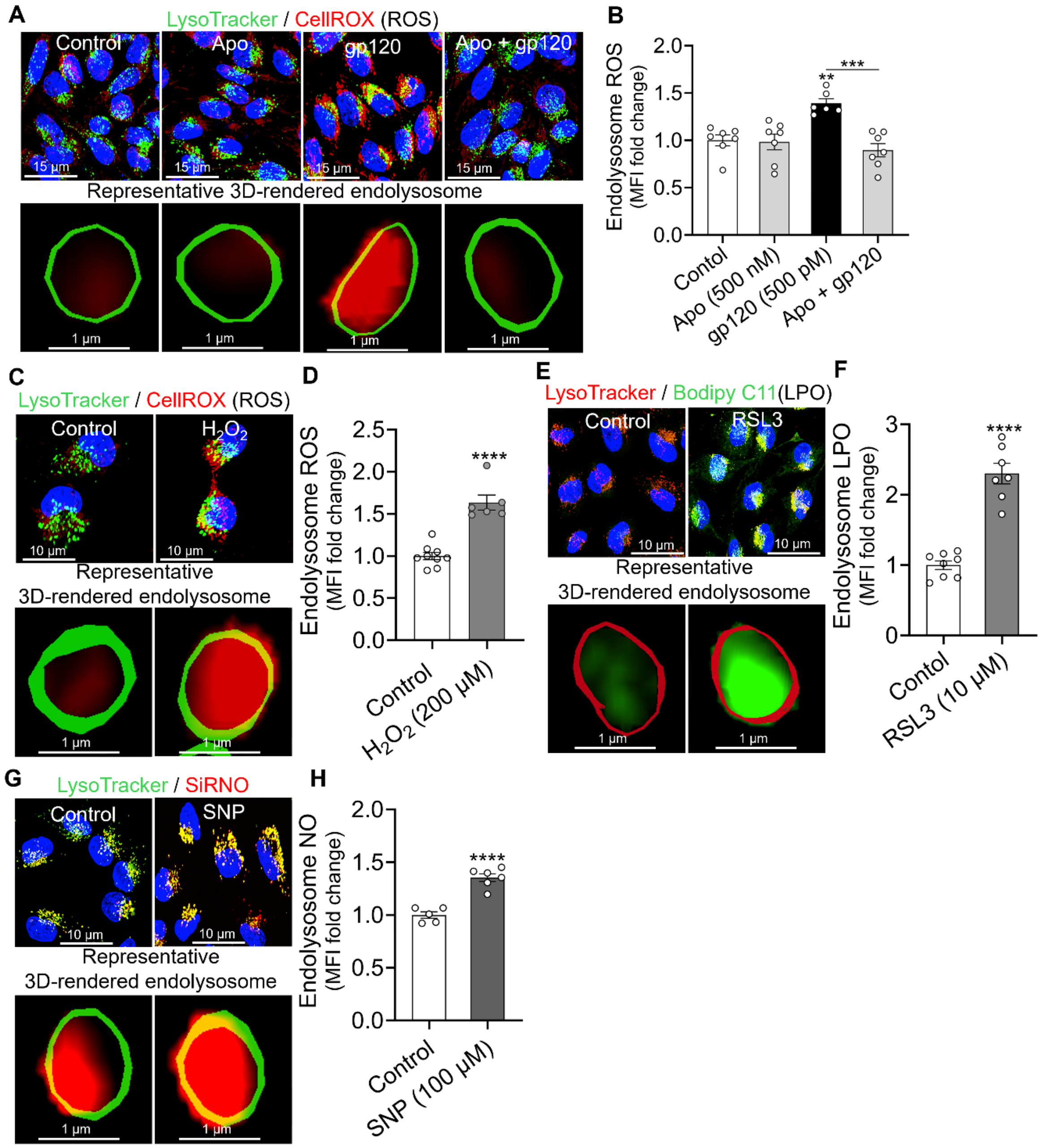


**Supplementary Fig. 2:** gp120-induced increases in endolysosome ROS were blocked by NOX2 inhibition, and different positive controls increased endolysosome ROS, lipid peroxidation and nitric oxide. **(A)** Representative confocal images of SH-SY5Y cells pretreated with the NOX2 inhibitor apocynin (500 nM, 30 min) or vehicle (control) and then treated with gp120 (500 pM, 4 h). Cells were stained with CellROX (red) for ROS, LysoTracker Green (green) for endolysosome, and Hoechst 33342 (blue) for nuclei. Scale bars = 10 µm. A representative

Imaris-generated surface rendering of a Lysotracker-positive endolysosome containing ROS fluorescence is also shown. Scale bar = 1 µm. **(B)** quantification of CellROX fluorescence in LysoTracker-positive endolysosomes. gp120-induced increases in endolysosome ROS were blocked by apocynin. **(C-E)** Representative confocal images of SH-SY5Y cells treated with $H_2O_2$ (200 µM, 4 h), RSL3 (10 µM, 4 h), SNP (100 µM, 4 h) or vehicle (control). Cells were stained with stained with CellROX (red) for ROS, Bodipy C11(green) for lipid peroxidation (LPO), or siRNO (red) for nitric oxide (NO), along with LysoTracker Green or Red for endolysosomes, and Hoechst 33342 (blue) for nuclei. Scale bars = 10 µm. Representative Imaris-generated Lysotracker-positive endolysosome containing ROS, LPO, or NO fluorescence are also shown. Scale bar = 1 µm. **(F-H)** Quantification of CellROX or Bodipy C11, or SiRNO fluorescence in LysoTracker-positive endolysosomes. $H_2O_2$ increased endolysosome ROS, RSL3 increased LPO, and SNP increased NO. Data are shown as mean ± SEM with individual data points (n = 5-9). Each data point represents MFI in LysoTracker-positive endolysosomes from at least 50 cells per condition. Statistical analysis was performed using Two-way ANOVA with Tukey's multiple comparison or two-tailed Student's *t*-test. $^{**}p < 0.01$, $^{***}p < 0.001$, $^{****}p < 0.0001$

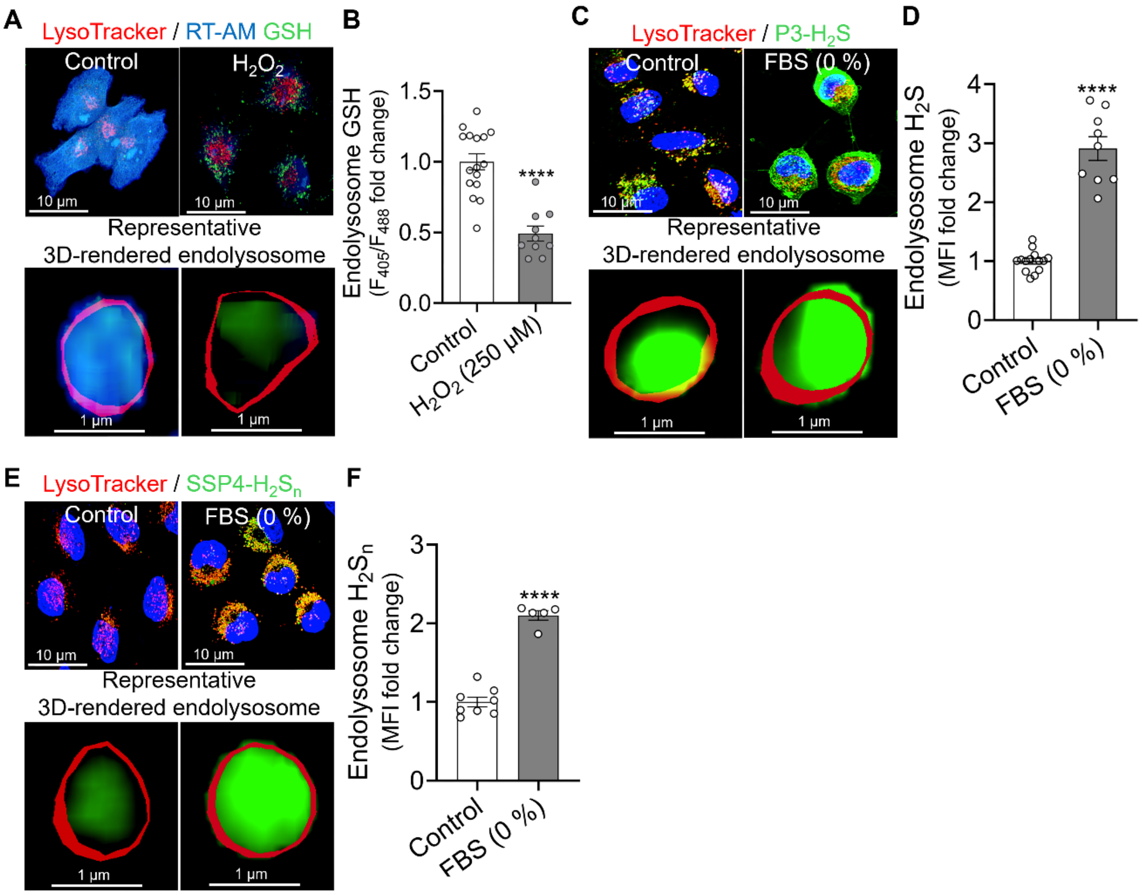


**Supplementary Fig. 3:** $H_2O_2$ decreases endolysosome GSH, and serum deprivation elevates $H_2S$ and sulfane sulfur. **(A,C,E)** Representative confocal images of SH-SY5Y cells treated with $H_2O_2$ (250 µM, 1 h) or vehicle control or incubated in FBS-free medium (0% FBS, 4 h) versus complete medium (control). Cells were stained with RT-AM GSH (blue/green) for GSH, P3 (green) for $H_2S$ or SSP4 (green) for sulfane sulfur ($H_2S_n$). Endolysosomes were labeled with LysoTracker Far Red (red) or LysoTracker Red, and nuclei with Hoechst 33342 (blue). Scale bars = 10 µm. A representative Imaris-generated surface rendering of a Lysotracker-positive endolysosome containing GSH , $H_2S$, or $H_2S_n$ fluorescence are also shown. Scale bar = 1 µm. **(B,D,F)** Measurement of RT-AM ratio (B), P3 (D), or SSP4 (F) fluorescence within LysoTracker-positive endolysosomes. Data are presented as fold change of mean fluorescence intensity (MFI). $H_2O_2$ significantly decreased endolysosome GSH, while serum deprivation increased endolysosome $H_2S$ and $H_2S_n$. Data are shown as mean ± SEM with individual data points (n = 5-16). Each data point represents the MFI within LysoTracker-positive endolysosomes from a minimum of 100 cells per condition. Two-tailed Student's *t*-test was used. ****$p < 0.0001$

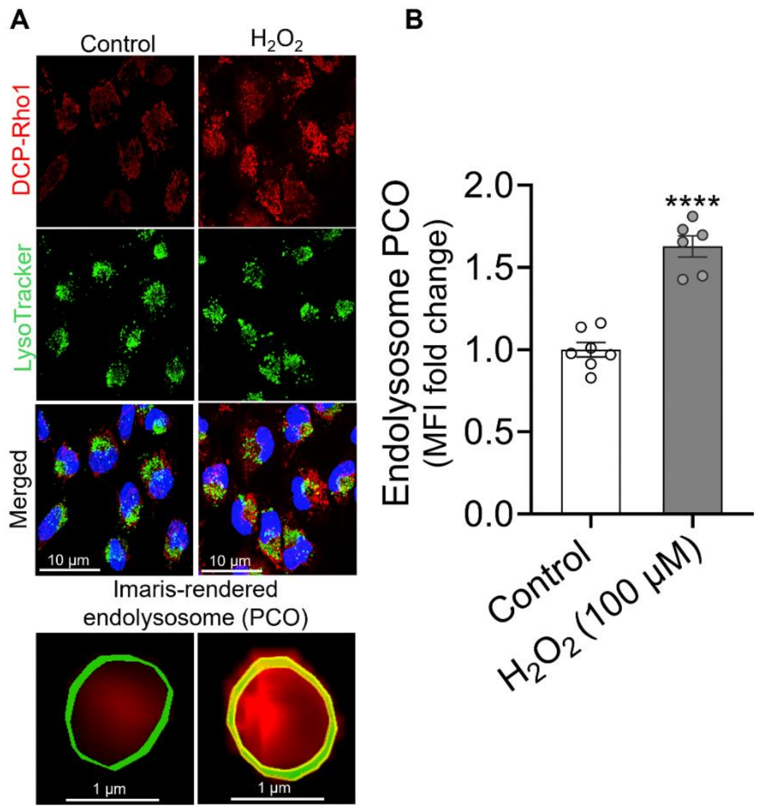


**Supplementary Fig. 4:** <u>$H_2O_2$ increases protein cysteine oxidation (PCO) in endolysosome proteins.</u> **(A)** Representative confocal images of SH-SY5Y cells treated with $H_2O_2$ (100 μM, 1 h) or vehicle (control) and stained with DCP-Rho1 (red) for protein cysteine oxidation (PCO), LysoTracker Green (green) for endolysosomes, and Hoechst 33342 (blue) for nuclei. Scale bars = 10 μm. A representative Imaris-generated surface rendering of a Lysotracker-positive endolysosome containing DCP-Rho1 fluorescence are also shown. Scale bar = 1 μm. **(B)** Measurement of DCP-Rho1 fluorescence within LysoTracker-positive endolysosomes using Imaris software. Data are shown as fold change of mean fluorescence intensity (MFI). $H_2O_2$ significantly increased endolysosome PCO. Data are presented as mean ± SEM with individual data points (n = 6-7). Each data point represents DCP-Rho1 MFI within LysoTracker-positive endolysosomes from a minimum of 100 cells per condition. Two-tailed Student's *t*-test was used for statistical analyses. *****p*< 0.0001

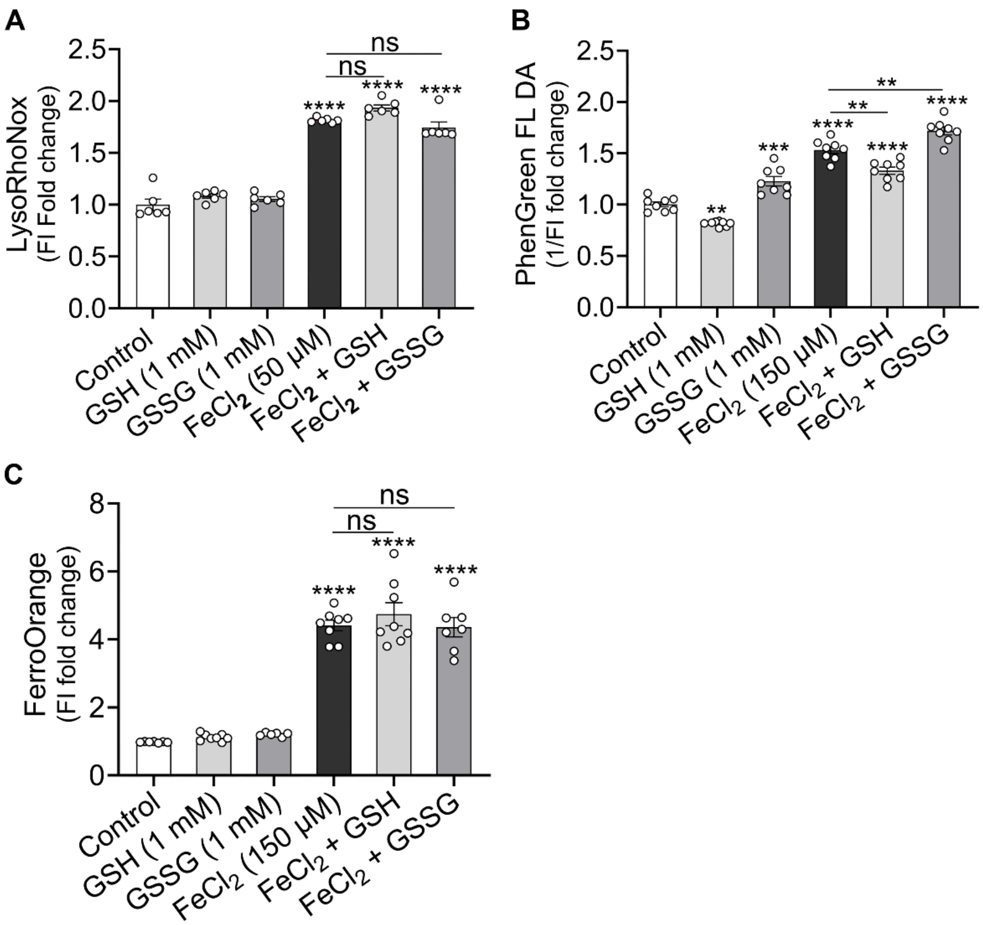


**Supplementary Fig. 5:** Effect of GSH and GSSG on $Fe^{2+}$ detection by fluorescent probes. LysoRhoNox (5 μM), PhenGreen FL DA (5 μM), or FerroOrange (1 μM) was diluted in PBS in 96-well plates, with or without GSH (1 mM) or GSSG (1 mM). $FeCl_2$ (50, or 150 μM) was then added, and fluorescence was measured using a microplate reader. Data are presented as fold change in fluorescence intensity. **(A)** LysoRhoNox fluorescence intensity (FI) was increased by $FeCl_2$ (50 μM) but not altered by GSH (1 mM) or GSSG (1 mM) alone or in combination with $FeCl_2$. **(B)** Reciprocal PhenGreen FI (1/FI) was increased by $FeCl_2$ (150 μM). GSH decreased whereas GSSG increased 1/FI values, both alone and with $FeCl_2$. **(C)** FerroOrange FI was increased by $FeCl_2$ (150 μM) but was unaffected by GSH or GSSG alone or in combination with $FeCl_2$. Data are shown as mean ± SEM with individual data points (n = 6-8). Each data point represents the fluorescence intensity from one well of a 96-well plate. Two-way ANOVA with Tukey's multiple comparison was used. ns = non-significant, $^{**}p<0.01$, $^{***}p<0.001$, $^{****}p<0.0001$
